# Quantitative mapping of cerebrovascular reactivity amplitude and delay with breath-hold BOLD fMRI when end-tidal CO_2_ quality is low

**DOI:** 10.1101/2024.11.18.624159

**Authors:** Rebecca G. Clements, Kristina M. Zvolanek, Neha A. Reddy, Kimberly J. Hemmerling, Roza G. Bayrak, Catie Chang, Molly G. Bright

## Abstract

Cerebrovascular reactivity (CVR), the ability of cerebral blood vessels to dilate or constrict in response to a vasoactive stimulus, is a clinically useful measure of cerebrovascular health. CVR is often measured using a breath-hold task to modulate blood CO_2_ levels during an fMRI scan. Measuring end-tidal CO_2_ (P_ET_CO_2_) with a nasal cannula during the task allows CVR amplitude to be calculated in standard units (vascular response per unit change in CO_2_, or %BOLD/mmHg) and CVR delay to be calculated in seconds. The use of standard units allows for normative CVR ranges to be established and for CVR comparisons to be made across subjects and scan sessions. Although breath holding can be successfully performed by diverse patient populations, obtaining accurate P_ET_CO_2_ measurements requires additional task compliance; specifically, participants must breathe exclusively through their nose and exhale immediately before and after each breath hold. Meeting these requirements is challenging, even in healthy participants, and this has limited the translational potential of breath-hold fMRI for CVR mapping. Previous work has focused on using alternative regressors such as respiration volume per time (RVT), derived from respiration-belt measurements, to map CVR. Because measuring RVT does not require additional task compliance from participants, it is a more feasible measure than P_ET_CO_2_. However, using RVT does not produce CVR amplitude in standard units. In this work, we explored how to achieve CVR amplitude maps, in standard units, and CVR delay maps, when breath-hold task P_ET_CO_2_ data quality is low. First, we evaluated whether RVT could be scaled to units of mmHg using a subset of P_ET_CO_2_ data of sufficiently high quality. Second, we explored whether a P_ET_CO_2_ timeseries predicted from RVT using deep learning allows for more accurate CVR measurements. Using a dense-mapping breath-hold fMRI dataset, we showed that both rescaled RVT and rescaled, predicted P_ET_CO_2_ can be used to produce maps of CVR amplitude in standard units and CVR delay with strong absolute agreement to ground-truth maps. The rescaled, predicted P_ET_CO_2_ regressor resulted in superior accuracy for both CVR amplitude and delay. In an individual with regions of increased CVR delay due to Moyamoya disease, the predicted P_ET_CO_2_ regressor also provided greater sensitivity to pathology than RVT. Ultimately, this work will increase the clinical applicability of CVR in populations exhibiting decreased task compliance.

## 1. Introduction

Maintaining appropriate cerebral blood flow is critical for supplying a sufficient stream of oxygen and nutrients to the brain. Cerebrovascular reactivity (CVR) reflects the ability of blood vessels in the brain to dilate or constrict in response to a vasoactive stimulus. CVR is typically measured using the dynamic response to a vasodilatory challenge and is complementary to steady-state measures like cerebral blood flow and cerebral blood volume (Liu et al., 2019). CVR has demonstrated clinical utility for a range of conditions including stroke (Papassin et al., 2021), carotid stenosis (Sobczyk et al., 2020), traumatic brain injury (Mathieu et al., 2020), Alzheimer’s disease (Yezhuvath et al., 2012), and multiple sclerosis (Chiarelli et al., 2022). CVR is also sensitive to healthy aging (Peng et al., 2018), cognitive function (D. Kim et al., 2021), and exercise (Murrell et al., 2013). In addition to being an important measure of vascular function, CVR has demonstrated potential for calibrating functional magnetic resonance imaging (fMRI) data in order to more confidently assess changes in neural activity (Davis et al., 1998; Liu et al., 2013).

While a range of imaging methods can be employed to measure CVR, blood-oxygenation level-dependent (BOLD) fMRI is most commonly used. BOLD signal contrast is affected by many factors including cerebral blood flow, cerebral blood volume, the metabolic rate of oxygen, and total hemoglobin levels (Kassner et al., 2010); however, signal changes are primarily driven by changes in cerebral blood flow (Mandell et al., 2008). An alternative to BOLD fMRI is dynamic susceptibility contrast (DSC)-MRI (Nighoghossian et al., 1997). DSC is less commonly used as it requires injection of a gadolinium-based contrast agent and is often paired with injection of a vasodilator such as acetazolamide; both preclude our ability to acquire repeated imaging measurements that would improve the robustness of CVR estimates. Arterial spin labeling (ASL)-MRI is another alternative that allows for noninvasive, direct quantification of cerebral blood flow. However, ASL has lower signal-to-noise ratio and poor temporal resolution as compared with BOLD fMRI techniques and is additionally biased by changes in labeling efficiency during changes in blood flow velocity (Pinto et al., 2021). Several review articles have discussed the utility of BOLD fMRI and alternative strategies for CVR mapping (Catchlove et al., 2018, Leoni et al., 2012, Liu et al., 2019, Pinto et al., 2021).

Typically, during a BOLD fMRI scan, arterial CO_2_ levels are deliberately increased to induce systemic vasodilation and thus increase blood flow. End-tidal CO_2_ (P_ET_CO_2_) values, which act as a surrogate for arterial CO_2_, are then used to compute CVR in standard units (%BOLD/mmHg). One common approach for increasing arterial CO_2_ involves intermittently inhaling air with a fixed concentration of CO_2_ (Lu et al., 2014). Computerized approaches have also been developed to dynamically change the inspired gas partial pressures to allow for precise targeting of P_ET_CO_2_ (Slessarev et al., 2007; Wise et al., 2007). While these gas delivery approaches allow for robust, reliable CVR characterization (Leung et al., 2016; Sobczyk et al., 2021), they require equipment that is often expensive, time-consuming to set up, and uncomfortable for participants. One highly feasible alternative involves using resting-state fMRI to map CVR by exploiting natural variations in arterial CO_2_ due to changes in breathing rate and depth (Golestani et al., 2016; Liu et al., 2017). However, a limitation of this approach is that spontaneous breathing changes may not cause sufficient BOLD signal variation to reliably assess CVR (Pinto et al., 2021). This is supported by De Vis et al. (2018), who found that a hypercapnia stimulus of at least 2 mmHg above baseline P_ET_CO_2_ is necessary to effectively evaluate hemodynamic impairment in a group of participants with internal carotid artery occlusive disease.

Completing breath holds during an fMRI scan is a promising method for robustly mapping CVR by invoking large changes in arterial CO_2_ levels without external gas delivery (Bright & Murphy, 2013; Kastrup et al., 2001). In addition to requiring less equipment than gas delivery methods, breath holds can also increase participant comfort since they do not require the participant to wear a face mask within the head coil and can be stopped by the participant at any time (Bright & Murphy, 2013). Rather than a face mask, participants typically wear a nasal cannula during the scan so that P_ET_CO_2_ can be measured to approximate arterial CO_2_ and calculate CVR. Compared to the face mask, participants often report the nasal cannula to be more comfortable due to its minimal contact with the face, breathability, and smaller size, allowing for a better fit within the head coil.

When measuring CVR, and particularly when characterizing a transient or dynamic response such as the response to a breath hold, it is important to consider both the amplitude and timing of the blood flow response. Variations in CVR timing can arise from regional heterogeneities in arterial transit times and variations in local vasodilatory response dynamics (Stickland et al., 2021). One approach for modeling CVR that accounts for both the amplitude and delay of the response at each voxel is a lagged general linear model framework (Moia et al., 2020a; Stickland et al., 2021). In this framework, multiple shifted variants of the P_ET_CO_2_ regressor are used to model the BOLD response to the P_ET_CO_2_ regressor at each voxel. The shift that maximizes the full model R^2^ is used to calculate the CVR amplitude and is considered the hemodynamic delay. Accounting for hemodynamic delays not only improves the accuracy of CVR amplitude estimates but also provides a complementary measure of cerebrovascular health (Donahue et al., 2015; Stickland et al., 2021). Additionally, since this approach utilizes a P_ET_CO_2_ regressor recorded during the scan, CVR amplitude can be calculated in standard units (%BOLD/mmHg) and CVR delay can be calculated in seconds.

The use of standard units allows normative CVR ranges to be established and CVR comparisons to be made across subjects and scan sessions. However, one challenge with this approach is obtaining accurate P_ET_CO_2_ measurements, particularly during breath-hold protocols. There are two main requirements that participants must meet for the recorded P_ET_CO_2_ values to accurately approximate arterial CO_2_ changes associated with the breath-hold task. First, participants must exhale immediately before and after each breath hold (Bright & Murphy, 2013; Murphy et al., 2011). This is necessary because true end-tidal gas values are only achieved at the end of expirations. However, exhaling before a breath hold may make the breath hold more challenging (although the duration of the breath hold can be shortened accordingly) and exhaling at the end of a breath hold must override and slightly delay the instinctive urge to take recovery breaths. Second, because P_ET_CO_2_ is typically measured using a nasal cannula, participants must only breathe through their nose for the entire experiment. If the participant fails to meet these two requirements, the P_ET_CO_2_ regressor will have missing data and will otherwise be an inaccurate approximation of arterial CO_2_, which will likely result in an inaccurate CVR estimate. Collectively these requirements raise concerns about breath-hold CVR accuracy in pediatric populations and clinical populations such as those with dementia, in which fMRI task compliance is often lower. In fact, a recent study which used a breathing task to map CVR in a pediatric cohort observed age-related differences in task compliance within the cohort, with younger participants having less reliable P_ET_CO_2_ values (Stickland et al., 2021).

Respiration volume per time (RVT) is an alternative metric to P_ET_CO_2_, which captures changes in breathing rate and depth that likely drive the majority of changes in arterial CO_2_ during voluntary breathing modulations (Birn et al., 2006). During task-free resting-state breathing, temporal fluctuations in RVT have been found to be highly correlated with P_ET_CO_2_ and to explain similar spatial and temporal BOLD signal variance (Chang & Glover, 2009). RVT can be measured by recording changes in respiration effort using a pneumatic belt worn around the chest or abdomen. Because RVT does not require the participant to exhale before and after each breath hold nor to breathe through their nose, it is often easier to obtain a high-quality and complete RVT trace than a P_ET_CO_2_ trace. Additionally, since respiration belts are commonly included with many scanner set-ups (Zvolanek et al., 2023) and relatively comfortable to wear, recording RVT is immediately feasible for most settings and participants.

Previously, Zvolanek et al. (2023) found that when P_ET_CO_2_ data quality is sufficient, RVT can produce CVR amplitude and delay maps that are comparable to those from P_ET_CO_2_. The authors defined “sufficient” data as having greater than 50% power in the dominant frequency range of the breath-hold task. Furthermore, they found that when sufficient P_ET_CO_2_ recordings are not available, RVT can recover CVR amplitude and delay maps, as long as the participant attempted the breath-hold task (Zvolanek et al., 2023). However, because RVT is measured in arbitrary units, they noted that one limitation of this approach is that the CVR amplitude maps generated using RVT are not in the standard CVR units of %BOLD/mmHg. This means that the CVR amplitude maps can only be used to make relative comparisons between brain regions of a single subject from a single scan and cannot be appropriately compared across subjects or scan sessions.

An alternative approach is using respiration-belt recordings to predict P_ET_CO_2_ and then mapping CVR using the predicted P_ET_CO_2_ timeseries. This approach may better model the BOLD response to changes in arterial CO_2_ compared to using RVT alone. Agrawal et al. (2023) demonstrated the feasibility of predicting the complete CO_2_ pressure timeseries from respiration-belt recordings in resting-state data using deep learning. Their predicted CO_2_ pressure timeseries achieved a Pearson correlation of 0.946 ± 0.056 with the ground-truth CO_2_; they also derived P_ET_CO_2_ from the predicted CO_2_ timeseries and achieved a correlation of 0.512 ± 0.269 with the ground truth. The authors noted that they tried to predict P_ET_CO_2_ directly from RVT, but their model performed poorly. Similar to Zvolanek et al. (2023), the authors pointed out that since respiration recordings are in arbitrary units, they could only predict z-normalized CO_2_ timeseries (0 mean and a standard deviation of 1), which would not allow for CVR amplitude mapping in standard units. Furthermore, the authors exclusively trained and validated their model using resting-state data and did not extend to breath-hold data.

The goal of the current study is to develop a strategy for mapping CVR amplitude in *standard units* (%BOLD/mmHg) and CVR delay, in breath-hold BOLD fMRI data, when P_ET_CO_2_ quality is low. In many cases, the participant performs all or most of the breath-hold trials in a session, but the P_ET_CO_2_ timeseries only shows an end-tidal CO_2_ increase for a subset of the trials. This often occurs when a participant does not successfully exhale after the breath-hold period or breathes through their mouth in certain trials. In these cases, we expect the RVT timeseries to show large decreases corresponding to all or most of the breath holds and the BOLD data to show signal increases, particularly in gray matter, during those same breath holds. Here, we propose to make RVT have units of mmHg by rescaling it have the same minimum and maximum as a reliable portion of high-quality measured P_ET_CO_2_ (i.e., one successfully completed breath-hold trial). Rescaling RVT to mmHg will allow CVR to be calculated in units of %BOLD/mmHg.

Next, we will investigate whether using a P_ET_CO_2_ regressor predicted from RVT using deep learning produces more accurate maps of CVR amplitude and delay than the rescaled RVT regressor. As mentioned, Agrawal et al. (2023) previously used deep learning to predict P_ET_CO_2_ from RVT in resting-state data but found that their model performed poorly; we hypothesize that since breath holds evoke larger fluctuations in P_ET_CO_2_ than resting-state, breath-hold data will allow for more robust predictions of P_ET_CO_2_ than in their original work. Since the magnitude of arterial CO_2_ varies significantly both within and between healthy participants and depends on a variety of factors such as the time of day, metabolism, sleep, and diet (Crosby & Robbins, 2004), and RVT is recorded in arbitrary units and varies with changes in belt position and tightness, we will not use RVT to infer the magnitude of arterial CO_2_. Instead, we will rescale the predicted P_ET_CO_2_ regressor to mmHg using the same methods used to rescale RVT. We hypothesize that the rescaled, predicted P_ET_CO_2_ regressor will allow for more accurate maps of CVR amplitude (%BOLD/mmHg) and delay than the rescaled RVT regressor.

Ultimately, we will evaluate the use of rescaled RVT and rescaled, predicted P_ET_CO_2_ regressors for mapping CVR in a subset of the publicly available EuskalIBUR dataset (Moia et al., 2020b), which provides breath-hold fMRI data for a group of densely-sampled participants. This dataset will allow us to comprehensively evaluate these strategies, ultimately providing guidance on the most robust method for mapping CVR in diverse clinical populations.

## 2. Methods

### 2.1. Data

#### 2.1.1. In-house training dataset

To train a model to predict P_ET_CO_2_ and determine model hyperparameters, we compiled a large dataset of physiological recordings during various breath-holding protocols. This dataset is available on OSF at https://doi.org/10.17605/OSF.IO/Y5CK4 (Clements et al., 2024) and consists of 245 total datasets collected from 56 individuals (26 ± 4 years, 35 M) at Northwestern University under studies approved by the Northwestern University Institutional Review Board. Written, informed consent was obtained from all participants for being included in this study. Each dataset consisted of expired CO_2_ pressure (mmHg) and respiration effort (arbitrary units) simultaneously recorded during a breath-hold task; additional details on data collection methods are described in Appendix A.

This training dataset was collected using 4 different breath-hold tasks. All tasks had multiple breath-hold trials, each of which consisted of a period of paced breathing (always 3 seconds in, 3 seconds out) followed by a breath hold, an exhalation, and a recovery period (all of varied lengths across tasks). Some tasks also incorporated a period of rest before or after the trials. The timings of each task and the number of datasets collected using each task are summarized in Supplementary Table S1. Tasks 1 and 2 were acquired in the MRI scan environment and compiled from previous studies in our lab; the MRI data associated with Tasks 1 and 2 is not used in this study. Tasks 3 and 4 were acquired outside the MRI environment specifically for this project. To ensure consistency across data collection environments, all participants were in the supine position and viewed the task stimuli on a monitor using a mirror. Stimuli were presented using PsychoPy (Peirce, 2007). Tasks 3 and 4 were designed to improve the generalizability of our modeling to any breath-hold task by incorporating randomized task timings. For Task 3, the breath-hold durations were 10, 12, 14, 16, 18, and 20 seconds, with the order of these durations randomized without replacement for each participant. For Task 4, the paced breathing duration, breath-hold duration, and recovery duration were randomized with replacement for each participant from pre-defined ranges. Additionally, for each breath hold in Task 4, there was a 10% chance that the hold was skipped and replaced with a rest period, mimicking participants who fail to perform the trial (e.g., when falling asleep in the scanner).

#### 2.1.2. EuskalIBUR testing dataset

To evaluate our P_ET_CO_2_ prediction accuracy, as well as evaluate the performance of both rescaled RVT regressors and rescaled, predicted P_ET_CO_2_ regressors for mapping CVR in breath-hold fMRI data, we used the publicly available EuskalIBUR dataset that was acquired by researchers at a different institution. This breath-hold dataset consists of both physiological and MRI data.

10 participants (32 ± 6 years, 5M) completed 10 weekly MRI scan sessions each; every session included a breath-hold task during an fMRI scan. Data for 7 of the 10 participants can be found on OpenNeuro at doi:10.18112/openneuro.ds003192.v1.0.1 (Moia et al., 2020b). The total dataset size was 99 sessions due to a software malfunction during physiological data collection for subject 10, session 1. Multi-echo fMRI data were acquired with the following parameters: 340 scans, TR=1.5 seconds, TEs = 10.6/28.69/46.78/64.87/82.96 ms. For additional details about the fMRI data acquisition and breath-hold task, as well as the acquisition of single-band reference (SBRef) images, a T1-weighted MP2RAGE, and a T2-weighted Turbo Spin Echo image, readers are referred to Moia et al. (2021). Exhaled CO_2_ and respiration effort were measured during each fMRI scan; additional details on data collection methods are provided in Appendix A.

#### 2.1.3. Physiological data processing and evaluation

All CO_2_ and respiration-belt data in both the training and testing dataset were processed using in-house MATLAB code (MathWorks, Natick, MA, R2022b). For the CO_2_ data, a peak-detection algorithm identified end-tidal peaks. The results of the algorithm were manually verified, and the peaks were linearly interpolated to create P_ET_CO_2_ timeseries with the same frequency as the original CO_2_ data. P_ET_CO_2_ timeseries were rescaled from units of Volts to mmHg using instructions from the manufacturer of the gas analyzer. For the respiration-belt data, alternating minima and maxima were identified using a peak-detection algorithm, manually verified, and used to calculate respiration volume per time (RVT) based on the method described by Birn et al. (2006). The RVT estimations were linearly interpolated to create RVT timeseries with the same frequency as the original respiration-belt data. Since this method requires alternating minima and maxima, we accounted for having two consecutive minima due to exhales before and after the breath hold by only included minima before the hold (Zvolanek et al., 2023).

Next, we assessed P_ET_CO_2_ data quality to ensure that only high-quality breath holds were used to train the model and evaluate model predictions. To assess P_ET_CO_2_ quality, the P_ET_CO_2_ change induced by each breath hold was calculated; a large P_ET_CO_2_ change indicates a high-quality measurement since breath holding causes CO_2_ to accumulate in the blood (Tancredi & Hoge, 2013). A custom Python script was developed that identified the peaks in the raw CO_2_ timeseries that were immediately before and after each breath hold. Then, the change in CO_2_ induced by each breath hold was calculated as the difference in amplitude between the peaks in each pair. Among breath holds causing positive CO_2_ changes, the mean and standard deviation CO_2_ increase was calculated. Breath holds that caused a CO_2_ increase greater than the mean minus 1 standard deviation were classified as “high-quality.” This threshold was chosen with the aim of classifying the majority of breath holds that caused any CO_2_ increase as high-quality, while still excluding breath holds that caused CO_2_ increases substantially lower than average. These low CO_2_ increases were likely due to participants breathing through their mouth or not fully exhaling after the breath hold. After quality assessment, P_ET_CO_2_ and RVT timeseries were downsampled to 10 Hz.

Next, we needed to account for delays between P_ET_CO_2_ and RVT, related to measurement delays caused by factors such as sampling line lengths, as well as physiological delays between changes in respiratory volume and subsequent changes in arterial CO_2_. Therefore, each P_ET_CO_2_ dataset was shifted to maximize its negative cross-correlation with each RVT dataset. A negative correlation between P_ET_CO_2_ and RVT is expected because breath holds cause simultaneous increases in arterial CO_2_ and decreases in respiratory volume. Because we expected the measurement delay to be greater for P_ET_CO_2_ than RVT, we only allowed for negative shifts, meaning that P_ET_CO_2_ could only be shifted earlier in time. For all P_ET_CO_2_ recordings, the maximum allowable shift was 30 seconds. This maximum shift was identified through trial and error to ensure that all of the calculated shifts were not consistently at the maximum value. After each P_ET_CO_2_ timeseries was shifted, data were trimmed from the end of the corresponding RVT signal to ensure that it was the same length as the shifted P_ET_CO_2_ timeseries.

Lastly, all P_ET_CO_2_ and RVT recordings were z-normalized (i.e., zero mean and unit standard deviation). The RVT timeseries were z-normalized to avoid biasing the model, since RVT is derived from respiration effort data that are recorded in arbitrary units. P_ET_CO_2_ timeseries were also z-normalized because, as previously explained, the magnitude of arterial CO_2_ cannot be inferred from RVT alone, and the scaling between these data types will likely vary between individuals. We chose not to convolve RVT or P_ET_CO_2_ with response functions prior to training the model; our rationale is described in Appendix B.

#### 2.1.4. MRI pre-processing

MRI data from the EuskalIBUR dataset were pre-processed for each scan session in which the associated P_ET_CO_2_ trace had all high-quality breath holds and for 2 additional scan sessions in which the majority of breath holds were low-quality. Pre-processing was performed using custom scripts which follow the same key steps described in Zvolanek et al. (2023). Scripts are available at https://github.com/BrightLab-ANVIL/PreProc_BRAIN and utilize both FSL (Cox, 1996; Jenkinson et al., 2012) and AFNI (Cox, 1996) commands. In summary, motion realignment, brain extraction, optimal combination of the echoes using tedana (DuPre et al., 2021; Kundu et al., 2012, 2013), and distortion correction were performed on the fMRI data. The MP2RAGE was brain extracted and used to generate a gray matter mask which was transformed into functional space.

### 2.2. Experiments

#### 2.2.1. Prediction of P_ET_CO_2_ from RVT

To model P_ET_CO_2_ from RVT, we used a 1D fully convolutional network (FCN), which is a type of convolutional neural network that does not have any fully connected layers (Agrawal et al., 2023).

##### 2.2.1.1. Implementation details

We segmented our delay-corrected, z-normalized training datasets into smaller data segments containing only high-quality breath holds. This approach maximized the size of our training dataset because we did not need to exclude entire P_ET_CO_2_ recordings containing one or more poorly performed breath-hold trials. Instead, we excluded only the low-quality portions, making use of the high-quality segments from the same recording. Additionally, this method ensured that our model was generalizable to breath-hold tasks of varying lengths. High-quality breath holds were identified using the methods outlined in Section 2.1.3; the intentionally skipped breath holds from Task 4 were also classified as high-quality so that they could be included in the training dataset to increase the generalizability of our model. It is important to note that these skipped breath holds are distinct from low-quality breath holds because while low-quality breath holds have missing or erroneous P_ET_CO_2_ data, skipped breath holds do not imply incorrect data in the P_ET_CO_2_ trace.

Next, P_ET_CO_2_ data were separated into blocks, each containing 1 breath hold. To ensure we captured the CO_2_ build-up and recovery effectively, we included data from before and after the apneic period. To identify the data that could be included in each segment, we calculated the halfway point between the end of each breath hold and the start of the next breath hold. Each block contained data from one halfway point to the next. We also manually estimated the start and end period of each skipped breath hold from Task 4 to create skipped breath-hold blocks. For blocks containing the first or the last trial in a dataset, we included all remaining data at the start or end of the trace, respectively. Using these breath-hold blocks, each P_ET_CO_2_ dataset was randomly segmented into 1–4 different data segments of varying lengths; each segment contained 2 or more consecutive, high-quality breath-hold blocks. The corresponding data segments from the RVT timeseries were extracted to be inputted to the model.

The input to the FCN is structured as an N x 2 array, with the first column containing the z-normalized, N-long RVT trace, and the second column containing the subject ID encoded using one-hot encoding (Paszke et al., 2019) and padded with zeroes to be N samples long. Subject ID was inputted to the model to account for the fact that most participants contributed multiple datasets and to encourage the model to learn subject-specific differences between physiological timeseries. Upon being inputted to the model, the z-normalized RVT trace was further normalized using the *tanh* operator to ensure that all values were between -1 and 1 (Agrawal et al., 2023). The output predicted P_ET_CO_2_ is an N x 1 array.

##### 2.2.1.2. Model optimization

Model hyperparameters were identified using only the in-house training dataset; the EuskalIBUR testing dataset was not used for hyperparameter optimization to avoid overfitting. Using the in-house training dataset, we performed 5-fold cross validation to ensure a robust estimation of model performance for each possible hyperparameter combination. For each fold, 80% and 20% of the data segments were assigned to the training and validation sets, respectively. The hyperparameters that we considered are summarized in Figure 1 and explained in more detail in the following paragraphs.

**Figure 1.**
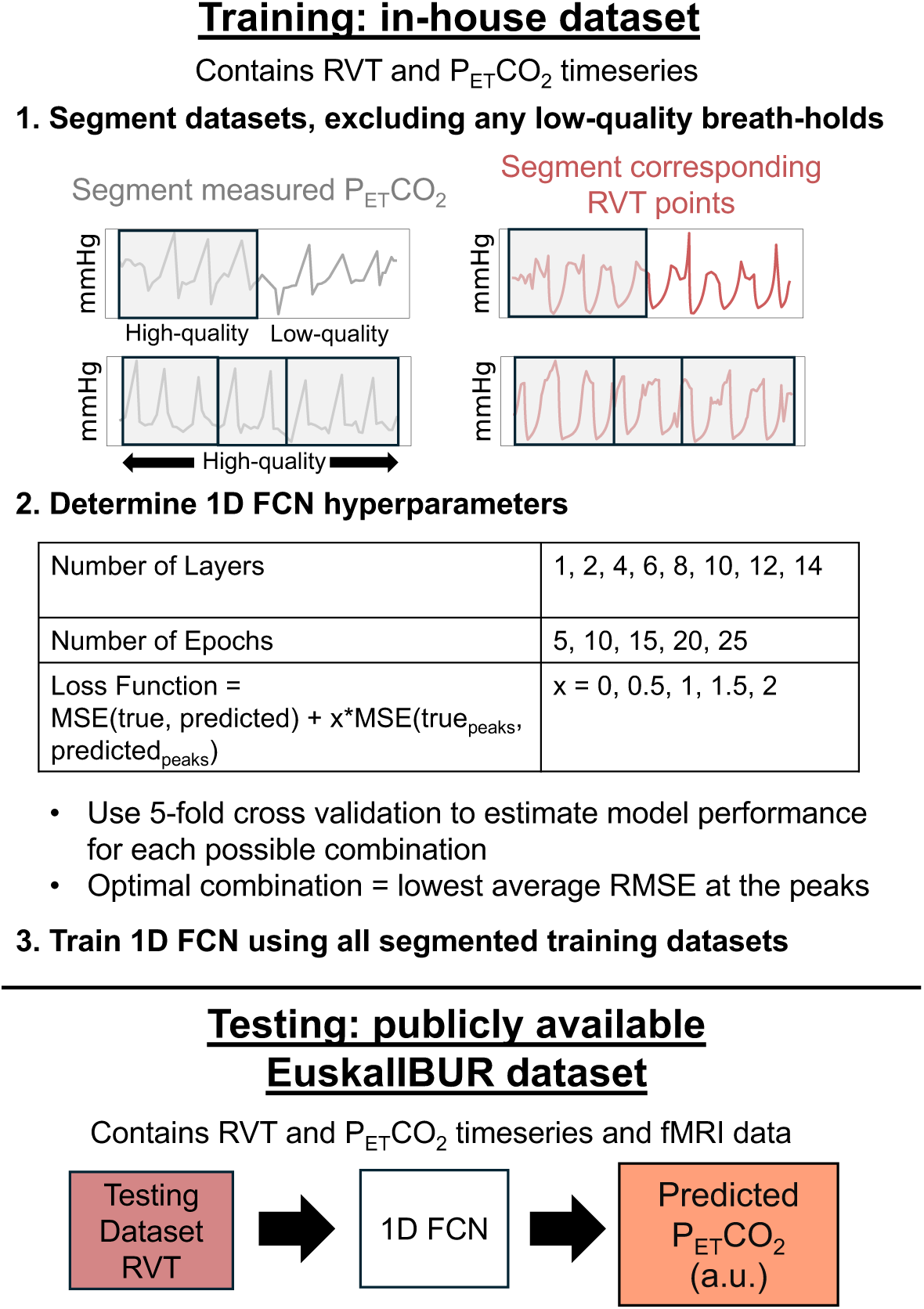
Overview of the methods for training and evaluating a 1D FCN to predict P_ET_CO_2_ from RVT. Training datasets were randomly separated into 1-4 smaller data segments, excluding any low-quality breath holds. Optimal hyperparameters were identified by using 5-fold cross-validation to estimate model performance for each possible combination, and the model was trained using the optimal hyperparameter combination. Next, the RVT timeseries in the test dataset, along with subject ID (not shown in the figure), were inputted to the model to generate P_ET_CO_2_ predictions in arbitrary units (a.u.).

First, we investigated using FCNs with varying numbers of hidden layers. Using methods described by Agrawal et al., who examined a similar relationship between physiological recordings in free (resting-state) breathing, we created FCNs with 1, 2, 4, and 6 convolutional layers. Because overfitting was not initially observed with the 6 convolutional layer model, we also created FCNs with 8, 10, 12, and 14 layers. For all models, half the layers were convolutional, and the other half were transposed convolutional. Both convolution and transposed convolution were performed using a stride of 2.

Each FCN used an adaptive learning rate that was implemented using Pytorch’s ReduceLROnPlateau command; the initial learning rate was 0.01, and this learning rate was reduced by a factor of 0.1 if improvements were not seen for 4 consecutive epochs (Paszke et al., 2019). Additionally, each FCN had a batch size of 1 and used the Adam optimization algorithm during training (Kingma & Lei Ba, 2015).

In addition to investigating different numbers of layers, we also investigated using 5, 10, 15, 20, and 25 epochs. For the loss function, we used mean squared error (MSE) between the measured and predicted P_ET_CO_2_ timeseries but, similarly to Agrawal et al. (2023), observed that when using MSE as the loss function, the FCN consistently underestimated the peaks in the data. In P_ET_CO_2_ timeseries, the values at the peaks are important since they indicate the extent of hypercapnia induced by each breath hold. Therefore, in addition to testing standard MSE, we tested four additional loss functions. These loss functions simply add the MSE at the peaks in the measured P_ET_CO_2_ trace, multiplied by a factor of either 0.5, 1, 1.5, or 2, to the standard MSE calculated using the entire P_ET_CO_2_ trace. Peaks were automatically identified using the *Peakutils* Python package.

For each possible hyperparameter combination, we compared the ground-truth and predicted P_ET_CO_2_ traces by calculating the average and standard deviation of the Pearson correlations transformed to Fisher’s Z across the 5 folds. We also calculated the average and standard deviation mean absolute error (MAE), root mean square error (RMSE), and RMSE at each of the peaks across the five folds. As described above, the height of the P_ET_CO_2_ peaks provides valuable information about the extent of hypercapnia during the breath hold, which is critical for accurately modeling CVR. Therefore, the hyperparameter combination that resulted in the lowest average RMSE at the peaks was considered optimal.

##### 2.2.1.3. Generation of predicted P_ET_CO_2_ timeseries

Using the optimal hyperparameters, we trained the FCN using the entire training dataset. After training was completed, the RVT timeseries and subject IDs for each subject in the testing dataset were inputted to the model to produce predicted P_ET_CO_2_ timeseries for each subject and session. A summary of the methods for training and optimizing the FCN and generating predicted P_ET_CO_2_ timeseries is provided in Figure 1.

#### 2.2.2. Rescaling of RVT and predicted P_ET_CO_2_ timeseries in the test set

Here, we investigated whether measured P_ET_CO_2_ data for one or more high-quality breath holds could be used to rescale RVT and predicted P_ET_CO_2_ to units of mmHg. Individual testing dataset breath-hold blocks, which included measured P_ET_CO_2_ data before, during, and after each high-quality breath hold, were used for rescaling. Breath hold blocks were generated using the same methods described in Section 2.2.1.1, except that for blocks containing the first trial in the dataset, we included enough data before the start of the first trial to make each block approximately the same length. Both predicted P_ET_CO_2_ and RVT were rescaled to have the same minimum and maximum as the first (high-quality) breath-hold block in the measured P_ET_CO_2_ timeseries. To better understand whether using more breath-hold blocks increased the rescaling accuracy, we also rescaled predicted P_ET_CO_2_ and RVT to have the same minimum and maximum as the first 2 sequential high-quality breath-hold blocks and the first 3 sequential high-quality breath-hold blocks. The result was three different sets of rescaled, predicted P_ET_CO_2_ and rescaled RVT regressors that were rescaled using high-quality measured P_ET_CO_2_ data from 1, 2, or 3 sequential breath-hold trials.

#### 2.2.3. Evaluation of rescaled RVT and rescaled, predicted P_ET_CO_2_ timeseries

Next, we assessed the error of rescaled RVT and rescaled, predicted P_ET_CO_2_ relative to measured P_ET_CO_2._ In these calculations, we only included datasets in which all of the breath holds in the measured P_ET_CO_2_ timeseries were classified as high-quality, meaning that these measured P_ET_CO_2_ timeseries could be used as ground truths. One caveat is that RVT is expected to be negatively correlated with measured P_ET_CO_2_, while predicted P_ET_CO_2_ is expected to be positively correlated with measured P_ET_CO_2_. To allow for fair comparisons and consistency with how these timeseries are typically processed in fMRI research, each measured and rescaled, predicted P_ET_CO_2_ timeseries was convolved with the canonical hemodynamic response function (HRF) (Friston et al., 1998) and each rescaled RVT timeseries was convolved with the respiration response function (RRF) (Birn et al., 2008). This made all of the timeseries positively correlated with each other, but the RRF and HRF have different latencies. Therefore, to evaluate rescaled RVT relative to measured P_ET_CO_2_, rescaled RVT timeseries convolved with the RRF were also shifted later in time (maximum shift = 30 seconds) to maximize their positive correlation with measured P_ET_CO_2_. To ensure that all signals being compared were the same length, measured and rescaled, predicted P_ET_CO_2_ timeseries, both convolved with the HRF, were trimmed to match the length of the shifted RVT timeseries convolved with the RRF.

To compare the strength of the relationships between RVT and measured P_ET_CO_2_, as well as between predicted P_ET_CO_2_ and measured P_ET_CO_2_, we calculated the mean and standard deviation Pearson’s correlation transformed to Fisher’s Z. Additionally, to evaluate the magnitude of differences between these metrics, we computed the average and standard deviation of the MAE, RMSE, and RMSE at the peaks of RVT and predicted P_ET_CO_2_ relative to measured P_ET_CO_2_ for each rescaling method.

Next, we conducted a 2-sided paired *t*-test (significance threshold *p*<0.05) to assess whether the correlations of RVT and predicted P_ET_CO_2_ to measured P_ET_CO_2_ were significantly different; only 1 *t*-test was needed since correlation is not affected by rescaling. For each error term (MAE, RMSE, and RMSE at the peaks), we conducted a repeated-measures 2-way ANOVA (significance threshold *p*<0.05) to test for an effect of the choice of regressor (RVT or predicted P_ET_CO_2_) and the number of breath holds used for rescaling (1, 2, or 3). Post hoc 2-sided paired *t*-tests were conducted to identify significantly different groups (significance threshold *p*<0.05, with Bonferroni correction).

To better understand our model’s P_ET_CO_2_ prediction performance, we also calculated the normalized correlation, MAE, RMSE, and RMSE at the peaks of measured and predicted P_ET_CO_2_ before either signal was convolved with the HRF, since convolution with the HRF can improve relationships between signals.

#### 2.2.4. Estimation of CVR amplitude and delay

For each pre-processed scan session in the EuskalIBUR dataset, 7 different regressors were used for 7 separate CVR calculations. These regressors were measured P_ET_CO_2_, predicted P_ET_CO_2_ rescaled using 1 breath hold, predicted P_ET_CO_2_ rescaled using 2 breath holds, predicted P_ET_CO_2_ rescaled using 3 breath holds, RVT rescaled using 1 breath hold, RVT rescaled using 2 breath holds, and RVT rescaled using 3 breath holds. Measured and predicted P_ET_CO_2_ regressors were convolved with the canonical HRF (Friston et al., 1998) and RVT regressors were convolved with the RRF (Birn et al., 2008). A summary of the methods used to generate these CVR regressors is shown in Figure 2. Voxelwise maps of CVR amplitude and delay were generated for each regressor with phys2cvr (Moia et al., 2024); additional details about this analysis are provided in Appendix C.

**Figure 2.**
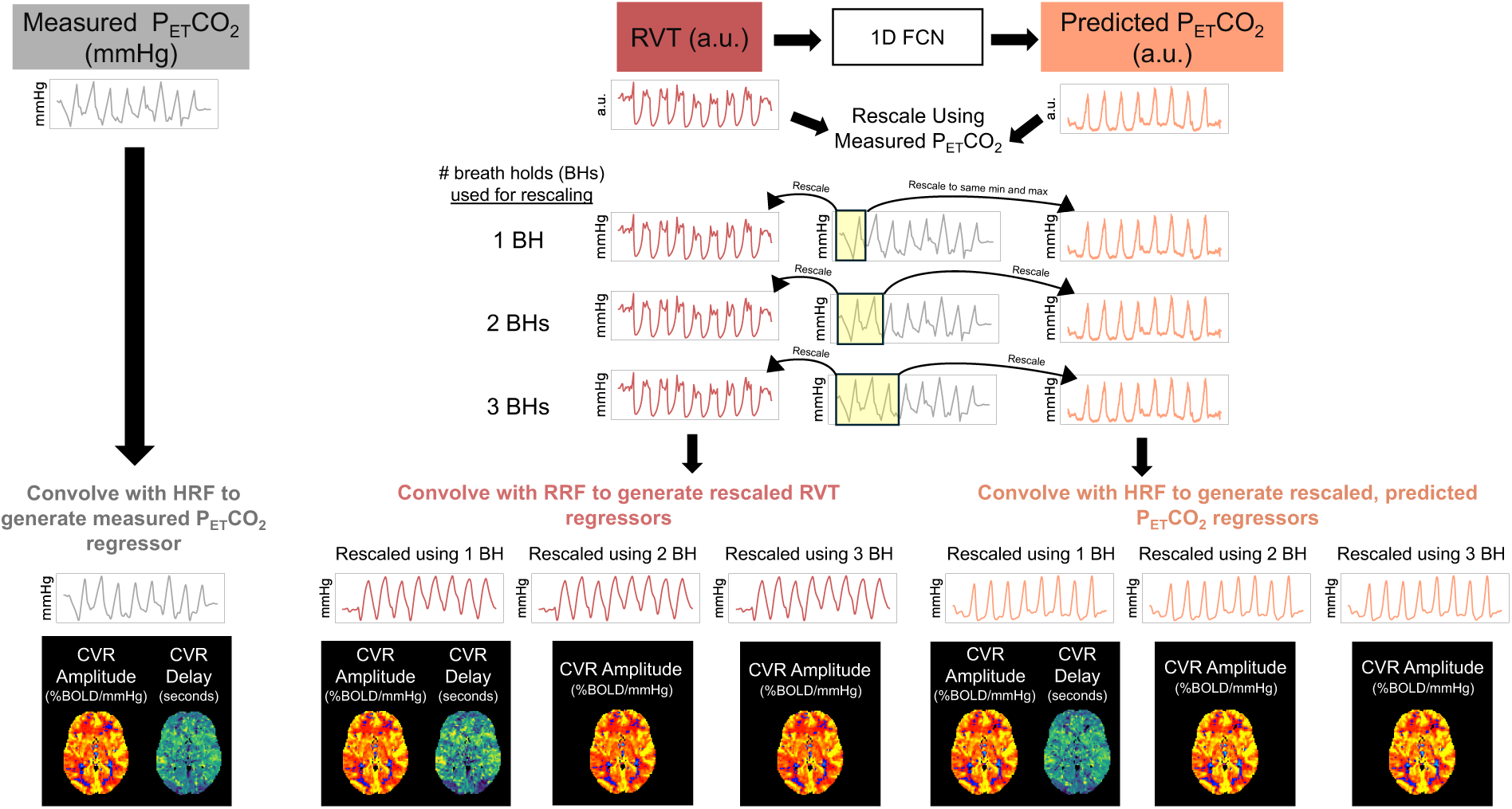
Overview of the methods pipeline for generating regressors to map CVR. Measured P_ET_CO_2_ was convolved with the canonical hemodynamic response function (HRF) and used to map CVR amplitude and delay. In datasets with P_ET_CO_2_ timeseries containing only high-quality breath holds, these maps served as ground truths. RVT was used as an input to a 1D FCN to generate predicted P_ET_CO_2_ timeseries in arbitrary units (a.u.). Both RVT and predicted P_ET_CO_2_ were rescaled to mmHg using 1, 2, and 3 breath holds in the measured P_ET_CO_2_ timeseries. Rescaled RVT and rescaled, predicted P_ET_CO_2_ were convolved with the respiration and hemodynamic response functions, respectively, and used to map CVR. Note that CVR delay is not sensitive to rescaling, so only 1 delay map each was generated for RVT and predicted P_ET_CO_2_.

#### 2.2.5. Analysis of CVR amplitude and delay maps for datasets with all high-quality breath holds

Datasets with measured P_ET_CO_2_ timeseries containing only high-quality breath holds provided CVR amplitude and delay maps that served as ground truths. Among these datasets, we calculated 6 group-level MAE maps to show CVR amplitude errors due to the choice of regressor (predicted P_ET_CO_2_ or RVT) and the choice of rescaling method (1, 2, or 3 breath holds) relative to the ground-truth maps. As CVR delay maps are not sensitive to the rescaling method, we also generated 2 MAE maps to show differences between delay maps generated using predicted P_ET_CO_2_ or RVT and ground-truth delay maps. For each MAE map, the median MAE in gray matter was calculated.

To better understand errors in scan-level CVR amplitude estimations related to the choice of regressor and the rescaling method, we also calculated the normalized correlation, MAE, and RMSE in gray matter for each individual CVR amplitude map relative to the ground-truth map. Voxels with CVR amplitude values above the 98^th^ percentile were excluded from these calculations. Since normalized correlation is not sensitive to rescaling, only 1 2-sided paired *t*-test (significance threshold *p*<0.05) was conducted to assess differences between the correlations of CVR amplitude values generated using RVT or predicted P_ET_CO_2_ to the ground-truth amplitude maps. A repeated-measures 2-way ANOVA (significance threshold *p*<0.05) was used to test for an effect of the choice of regressor (RVT or predicted P_ET_CO_2_) and the number of breath holds used for rescaling (1, 2, or 3) on the MAE and RMSE of CVR amplitude in gray matter, and post hoc 2-sided paired *t*-tests were conducted to identify significantly different groups (significance threshold *p*<0.05, with Bonferroni correction).

Lastly, we assessed how well the predicted P_ET_CO_2_ and RVT regressors maintain the ranking of CVR amplitude values across subjects and scan sessions. Specifically, we wanted to confirm that if a subject exhibited a particularly high or low ground-truth CVR value for a particular scan compared to all other scans, this subject would also show a relatively high or low CVR value in maps generated using the rescaled, predicted P_ET_CO_2_ and rescaled RVT regressors. To assess this, the Spearman Rank Correlation was calculated to compare the rankings of median CVR amplitudes in gray matter for measured P_ET_CO_2_ (ground-truth) and RVT amplitude maps, as well as measured P_ET_CO_2_ and predicted P_ET_CO_2_ amplitude maps. This analysis was performed for RVT and predicted P_ET_CO_2_ regressors rescaled using 1, 2, and 3 breath holds.

#### 2.2.6. Analysis of CVR amplitude and delay maps in datasets with mostly low-quality breath holds

We also evaluated the effectiveness of using a rescaled RVT and rescaled, predicted P_ET_CO_2_ regressor in 2 example datasets in which the RVT timeseries indicated that the participant attempted every breath hold in the task, but the P_ET_CO_2_ timeseries contained mostly low-quality breath holds (likely due to the participant not exhaling immediately after each breath hold). For each CVR amplitude and delay map generated using RVT and predicted P_ET_CO_2_, we calculated the spatial correlation in gray matter (3ddot, AFNI) relative to the ground-truth maps. For each scan, the ground-truth maps were from a different session for the same subject that had a measured P_ET_CO_2_ trace containing only high-quality breath holds.

#### 2.2.7. Case study in a participant with Moyamoya disease

To better understand the utility of using rescaled RVT and rescaled, predicted P_ET_CO_2_ regressors to map CVR, we also scanned a 31-year-old male with unilateral Moyamoya disease causing an occluded right middle cerebral artery. This dataset was collected at Northwestern University under a study approved by the Northwestern University Institutional Review Board; written, informed consent was obtained from the participant for being included in this study. The participant completed a breath-hold task during a functional T2*-weighted scan which used a multi-echo, gradient-echo EPI sequence (CMRR, Minnesota) on a 3T Siemens Prisma with a TR of 1.5 seconds. The TR was the same as the EuskalIBUR dataset, and the other functional scan parameters and breath-hold task were also similar to those in the EuskalIBUR dataset (Moia et al., 2020b, 2021). A whole brain T1-weighted EPI-navigated multi-echo MPRAGE scan, based on Tisdall et al. (2016), was also acquired with scan parameters previously described by Stickland et al. (2021).

Exhaled CO_2_ and respiration effort timeseries were recorded and processed (see Appendix A and Section 2.1.3, respectively, for methods), and predicted P_ET_CO_2_ timeseries were generated (Section 2.2.1.3). Predicted P_ET_CO_2_ and RVT were rescaled using measured P_ET_CO_2_ data for 1 high-quality breath hold (Section 2.2.2). fMRI data were pre-processed using similar methods as those described in Section 2.1.4. CVR amplitude and delay maps were calculated for the measured P_ET_CO_2_, rescaled, predicted P_ET_CO_2_, and rescaled RVT regressors using the methods described in Appendix C, with two exceptions to account for the expected increase in CVR delays due to Moyamoya pathology: a maximum lag value of ±15 seconds was used, and maps were not thresholded to remove voxels with delay values at the boundaries (-15, -14.7, 14.7, 15).

For both CVR amplitude and delay, within a gray matter mask, we evaluated the spatial correlation of the maps generated using rescaled, predicted P_ET_CO_2_ and rescaled RVT relative to the ground-truth amplitude and delay maps generated using measured P_ET_CO_2_ (3ddot, AFNI). Because we expected longer CVR delays in the right MCA territory in this participant (Stickland et al., 2021), we also specifically assessed whether the RVT and predicted P_ET_CO_2_ methods could be used to effectively identify brain areas with extreme delays. To identify these areas, we thresholded the delay maps to only contain voxels with delay values greater than or equal to 10 seconds and then made a binarized map of clusters with at least 15 voxels (3dClusterize, AFNI). Then, we calculated the Dice similarity coefficient between the clusters in the rescaled RVT delay map or the rescaled, predicted P_ET_CO_2_ delay map and the clusters in the measured P_ET_CO_2_ map (the ground truth).

## 3. Results

### 3.1. Physiological data processing and evaluation

In the in-house training dataset, the average CO_2_ change across all breath holds was 9.85 ± 3.51 mmHg. Any breath hold that resulted in a CO_2_ increase greater than 6.33 mmHg (the mean CO_2_ increase minus 1 standard deviation) was considered high-quality. In the EuskalIBUR test dataset, the average CO_2_ increase induced by a breath hold was 6.73 ± 3.13 mmHg and high-quality breath-hold trials needed to cause a CO_2_ increase greater than 3.60 mmHg.

We observed similar task compliance trends in the training and test datasets. In the training dataset, 81% of the 1708 individual breath-hold trials were high-quality. Additionally, 53% of the 245 total CO_2_ recordings contained all high-quality breath holds, and 4.5% of the CO_2_ recordings contained no high-quality breath holds. In the test dataset, 73% of the 800 individual trials were high-quality. 57% of the 99 total CO_2_ recordings contained entirely high-quality breath-hold trials, while 7% contained no high-quality breath-hold trials. In the test dataset, we observed that datasets with all high-quality breath-hold trials tended to have higher correlations between measured P_ET_CO_2_ and RVT than datasets with one or more low-quality trials (Supplementary Table S2)

Each P_ET_CO_2_ timeseries was shifted to account for delays between P_ET_CO_2_ and RVT. In the training and test datasets, P_ET_CO_2_ was shifted an average of 16.0 ± 3.6 and 23.5 ± 4.6 seconds earlier, respectively, to maximize its negative cross-correlation with RVT. Using the temporal location of each high-quality breath hold in each P_ET_CO_2_ timeseries in the training dataset (after accounting for the applied temporal shift), delay-corrected P_ET_CO_2_ and RVT timeseries in the training dataset were randomly segmented into 1-4 data segments consisting of consecutive, high-quality breath holds. After segmentation, the final size of the training dataset was 340 sets of P_ET_CO_2_ and RVT segments from 54 unique participants.

### 3.2. Model optimization

The model which resulted in the lowest RMSE at the peaks, and consequently was considered the optimal model, used 12 layers, 20 epochs, and a loss function which summed the standard MSE (calculated using all datapoints) with the MSE at the peaks scaled by 0.5. Averaged across all 5 folds, this model resulted in a mean Fisher’s Z of 1.299 ± 0.216, an MAE of 0.447 ± 0.111 (a.u.), an RMSE of 0.601 ± 0.124 (a.u.), and an RMSE at the peaks of 0.614 ± 0.486 (a.u.). Hyperparameters and error terms for the 5 next best performing models are provided in Supplementary Table S3.

### 3.3. Evaluation of rescaled RVT and rescaled, predicted P_ET_CO_2_ timeseries

The EuskalIBUR testing dataset (unused during optimization and training of the P_ET_CO_2_ prediction model) was used to evaluate the rescaled RVT and rescaled, predicted P_ET_CO_2_ timeseries relative to measured P_ET_CO_2_. Figure 3 shows example measured P_ET_CO_2_ timeseries plotted against predicted P_ET_CO_2_ and RVT timeseries, both of which were rescaled using 1 breath hold. Examples are provided for both measured P_ET_CO_2_ timeseries with only high-quality breath holds (Figure 3A) and measured P_ET_CO_2_ timeseries with mostly low-quality breath holds (Figure 3B).

**Figure 3.**
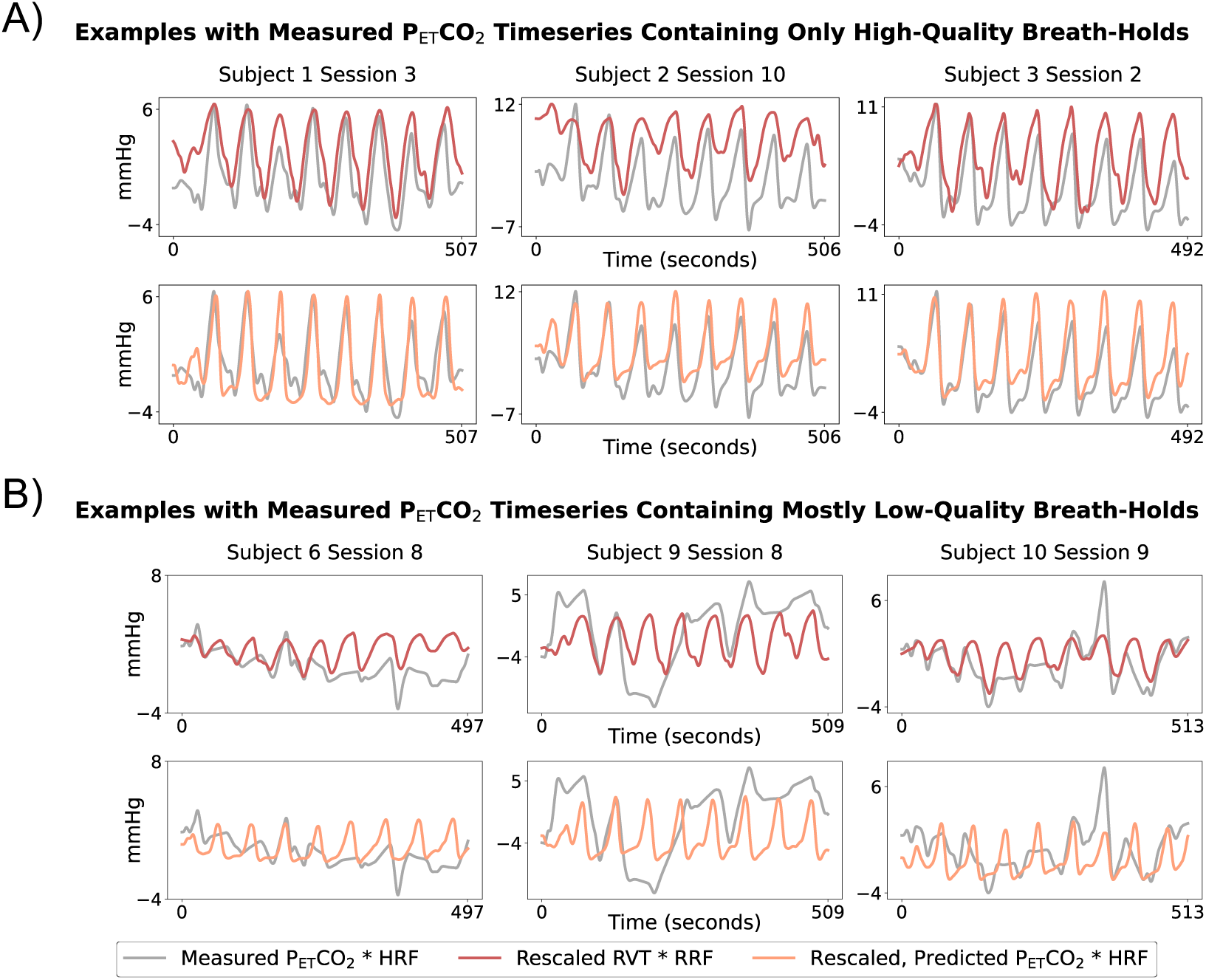
Example rescaled, predicted P_ET_CO_2_ and rescaled RVT timeseries (rescaled using 1 breath hold) plotted against measured P_ET_CO_2_ timeseries. Measured and predicted P_ET_CO_2_ timeseries are shown after convolution with the HRF, and RVT timeseries are shown after convolution with the RRF. Results are shown for 3 datasets in which the measured P_ET_CO_2_ had all high-quality breath holds (A) and 3 datasets with multiple low-quality breath holds (B).

Only datasets with measured P_ET_CO_2_ timeseries containing all high-quality breath holds were considered to be reasonable ground truths and were included in the following analysis (N=56). We compared rescaled RVT and rescaled, predicted P_ET_CO_2_ timeseries to measured P_ET_CO_2_ using a Pearson correlation normalized to Fisher’s Z, MAE, RMSE, and RMSE at the peaks (Figure 4). Predicted P_ET_CO_2_ had a significantly higher correlation with measured P_ET_CO_2_ than RVT (*p*<0.05). Our repeated-measures 2-way ANOVA showed that the choice of regressor and the number of breath holds used for rescaling had statistically significant main effects on the MAE and RMSE but not the RMSE at the peaks. Post hoc tests showed that across the 3 rescaling methods, RVT had a significantly higher MAE and RMSE than predicted P_ET_CO_2_ (*p*<0.05, Bonferroni corrected). For rescaling predicted P_ET_CO_2_, using 2 breath holds compared to 1 breath hold significantly decreased the MAE (*p*<0.05, Bonferroni corrected) but not the RMSE; using 3 breath holds compared to 2 did not significantly impact the MAE or RMSE. For RVT, rescaling using 2 breath holds compared to 1 breath hold did not significantly change the MAE or RMSE; rescaling using 3 breath holds compared to 2 breath holds significantly decreased the RMSE and MAE. Effect sizes and *p*-values for all comparisons are shown in Supplementary Table S4.

**Figure 4.**
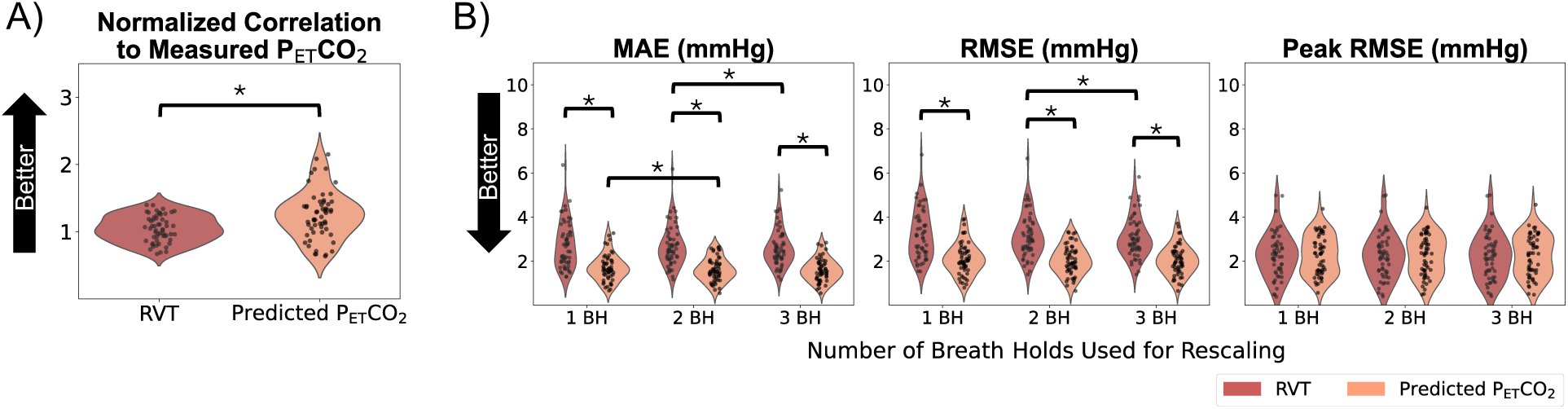
Overview of metrics comparing rescaled, predicted P_ET_CO_2_ and rescaled RVT to measured P_ET_CO_2_ in datasets in which all breath holds in the measured P_ET_CO_2_ timeseries were classified as high-quality. Both Fisher’s Z values (A), which are not affected by rescaling, and error terms for each rescaling method (B) are shown. Asterisks indicate significant differences.

To gain a better understanding of our model’s performance, we also assessed the correlation and error of predicted P_ET_CO_2_ relative to measured P_ET_CO_2_ before either signal was convolved with the HRF (Supplementary Figure S1). As expected, the mean normalized correlation of the unconvolved timeseries was slightly lower than that of the convolved timeseries (1.14 ± 0.32 compared to 1.24 ± 0.36); however, a normalized correlation of 1.14 still indicates strong P_ET_CO_2_ prediction performance. The error terms before and after convolution were relatively similar. These results suggest that our P_ET_CO_2_ prediction method is not restricted to the canonical HRF and that any appropriate response function can be effectively utilized with this approach.

### 3.4. CVR amplitude and delay maps for scans with all high-quality breath holds

For scan sessions with P_ET_CO_2_ timeseries containing all high-quality breath holds, CVR amplitude maps (in %BOLD/mmHg) generated using RVT and predicted P_ET_CO_2_ regressors, rescaled using 1 breath hold, appear similar (i.e., show similar amplitude patterns across the brain and have similar amplitude magnitudes) to the ground-truth maps generated using the measured P_ET_CO_2_ regressors (Figure 5). Similarly, the associated rescaled RVT and rescaled, predicted P_ET_CO_2_ CVR delay maps (in seconds, normalized to median gray matter delay) appear similar to the ground-truth measured P_ET_CO_2_ CVR delay maps (Figure 6). Difference maps showing the error of CVR amplitude and delay maps generated using rescaled RVT and rescaled, predicted P_ET_CO_2_ regressors for 8 example subjects are shown in Supplementary Figures S2 and S3. The delay maps generated using rescaled, predicted P_ET_CO_2_ seem to better estimate extreme negative or positive delay values (voxels that are yellow or dark purple) in the ground-truth delay map than the rescaled RVT maps. This is particularly evident in the delay maps for subject 10 session 4. Additionally, the delay maps generated using RVT appear to introduce extreme delay values that are not present in the ground-truth maps, as seen in the left posterior portion of the delay map for subject 3 session 2.

**Figure 5.**
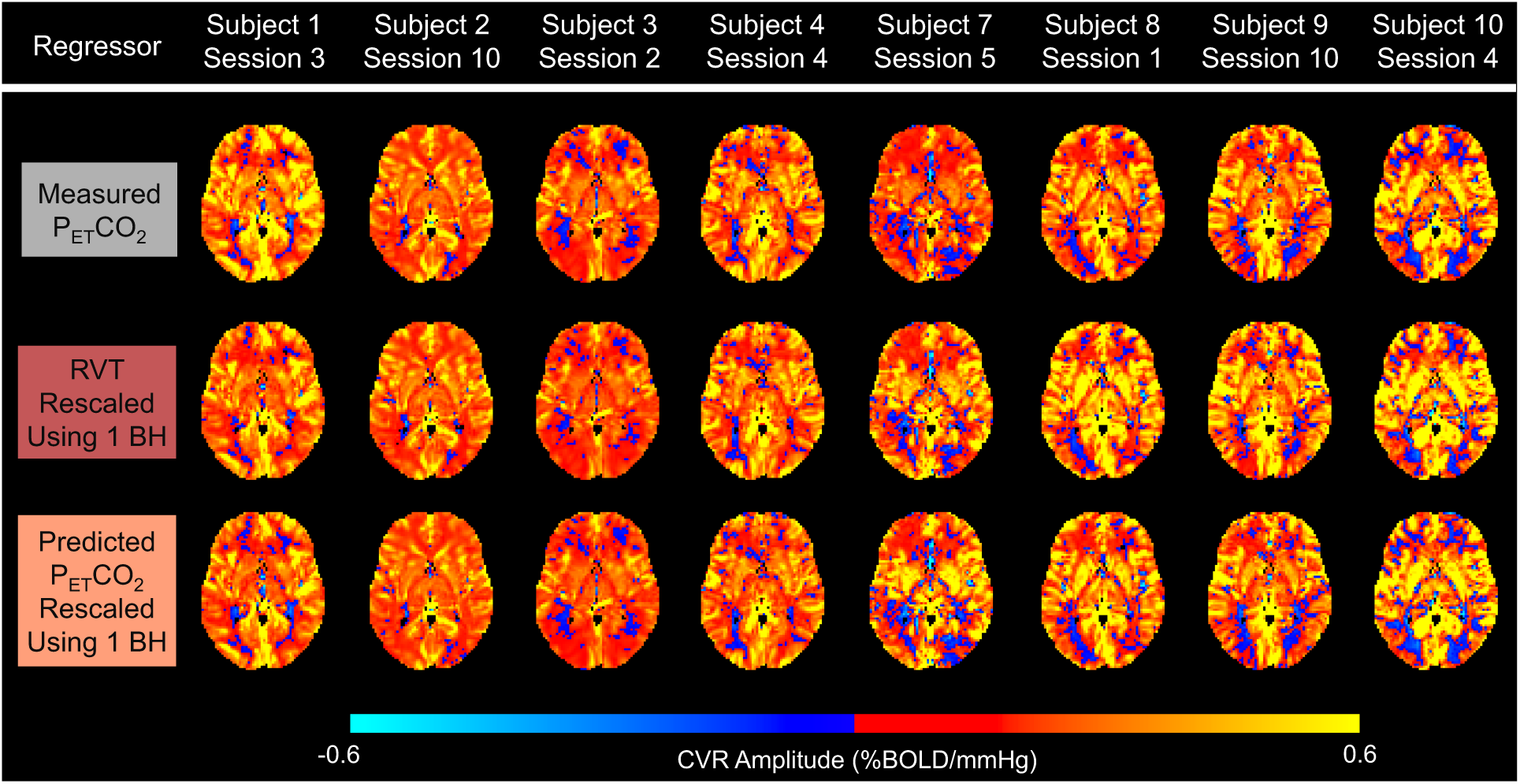
Example CVR amplitude maps for 8 subjects for sessions with measured P_ET_CO_2_ timeseries containing all high-quality breath holds (BHs). The top row shows ground-truth amplitude maps, generated using the measured P_ET_CO_2_ regressor, while the middle and bottom rows show maps generated using rescaled RVT and rescaled, predicted P_ET_CO_2_ regressors, respectively.

**Figure 6.**
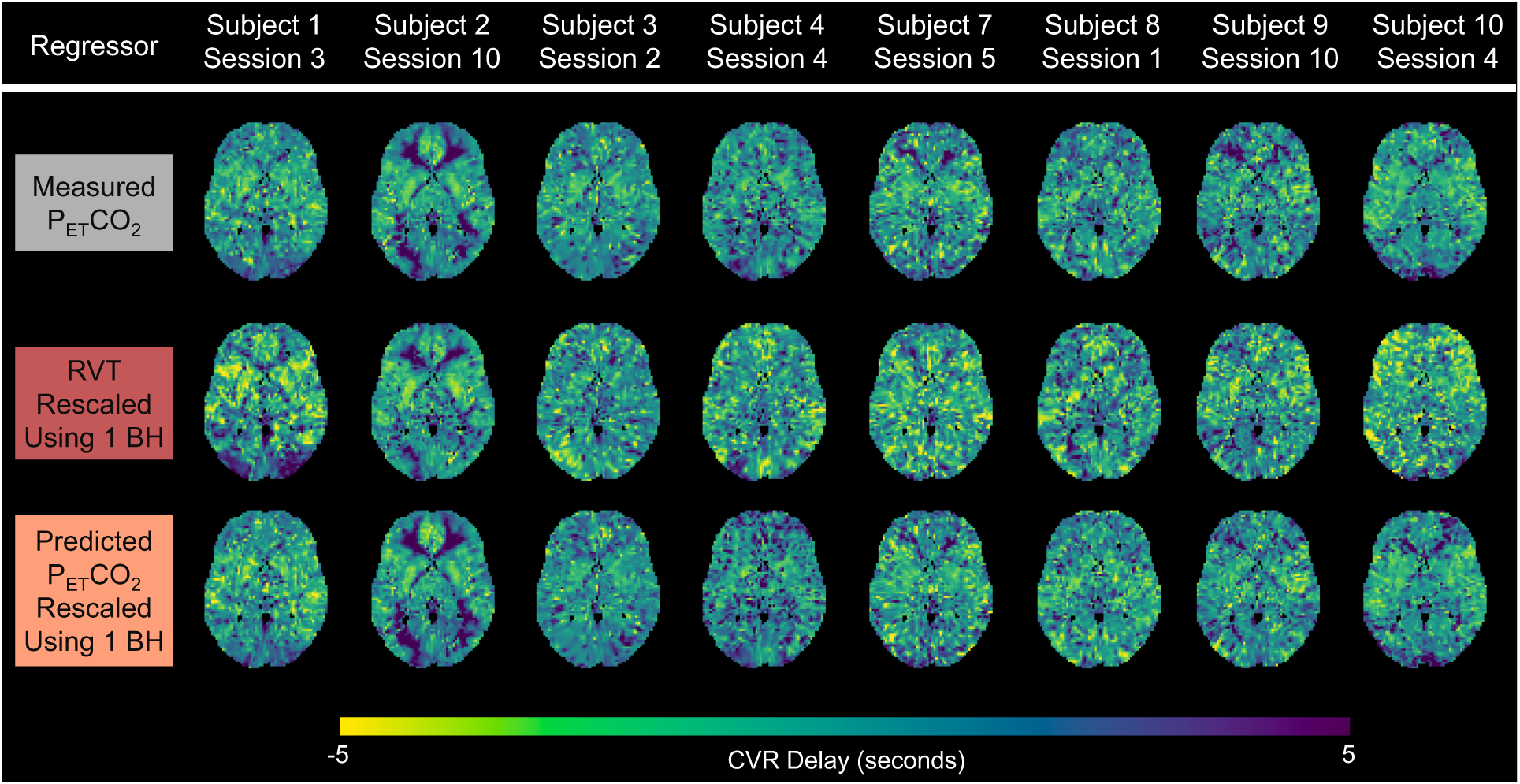
Example CVR delay maps for 8 subjects for sessions with measured P_ET_CO_2_ timeseries containing all high-quality breath holds (BHs). Ground-truth delay maps, generated using the measured P_ET_CO_2_ regressor, are shown in the top row, while delay maps generated using rescaled RVT and rescaled, predicted P_ET_CO_2_ regressors are shown in the middle and bottom rows, respectively. Delay maps are normalized to the gray matter median. Negative delays reflect earlier responses, while positive delays reflect later responses.

Next, to better understand the impact of the rescaling method on CVR accuracy, group-level MAE maps were computed to assess the errors in CVR amplitude maps generated using RVT and predicted P_ET_CO_2_ regressors, rescaled using 1, 2, and 3 breath holds, relative to the ground-truth maps generated using measured P_ET_CO_2_ (Figure 7). Across the 3 rescaling methods, the median MAE in gray matter was consistently lower for rescaled, predicted P_ET_CO_2_ than for RVT. In each map, the magnitude of the error appears consistent throughout gray matter, suggesting that the CVR amplitude bias introduced by the RVT and predicted P_ET_CO_2_ regressors is not specific to any part of the cortex. Increasing the number of breath holds used to rescale RVT slightly decreased the median MAE in gray matter. Rescaling predicted P_ET_CO_2_ using 2 breath holds compared to 1 but not 3 breath holds compared to 2 slightly decreased the median MAE in gray matter.

**Figure 7.**
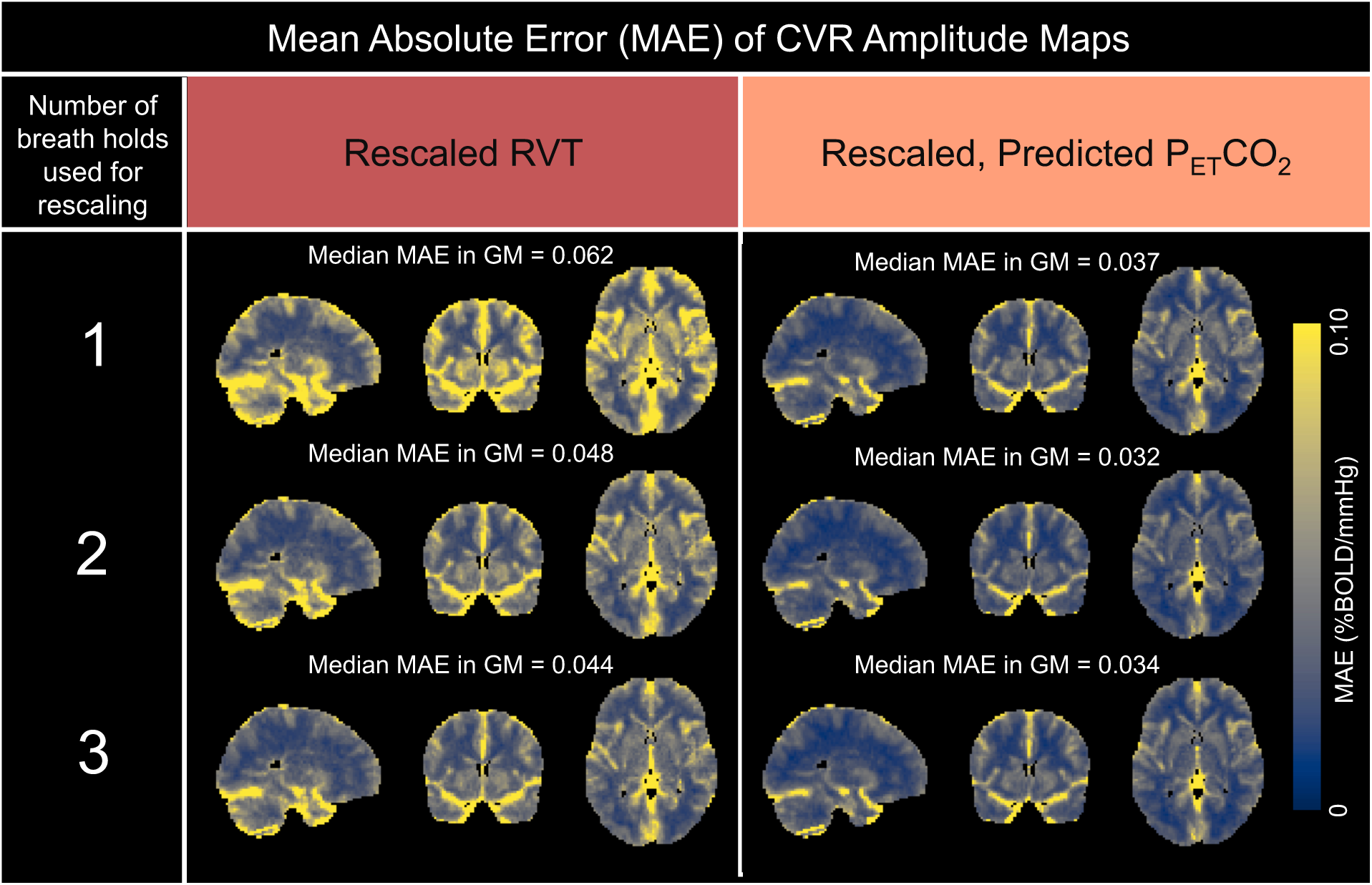
MAE maps comparing amplitude values generated using rescaled RVT and rescaled, predicted P_ET_CO_2_ to the ground-truth amplitude values generated using measured P_ET_CO_2_. Maps are shown for each of the 3 different rescaling methods. The median MAE in gray matter (%BOLD/mmHg) is shown above each map.

In terms of CVR delay, which is not sensitive to rescaling, the predicted P_ET_CO_2_ method outperformed the RVT method, with a median MAE in gray matter of 0.97 seconds compared to 1.51 seconds (Figure 8). Again, the errors for both maps appear relatively consistent throughout gray matter.

**Figure 8.**
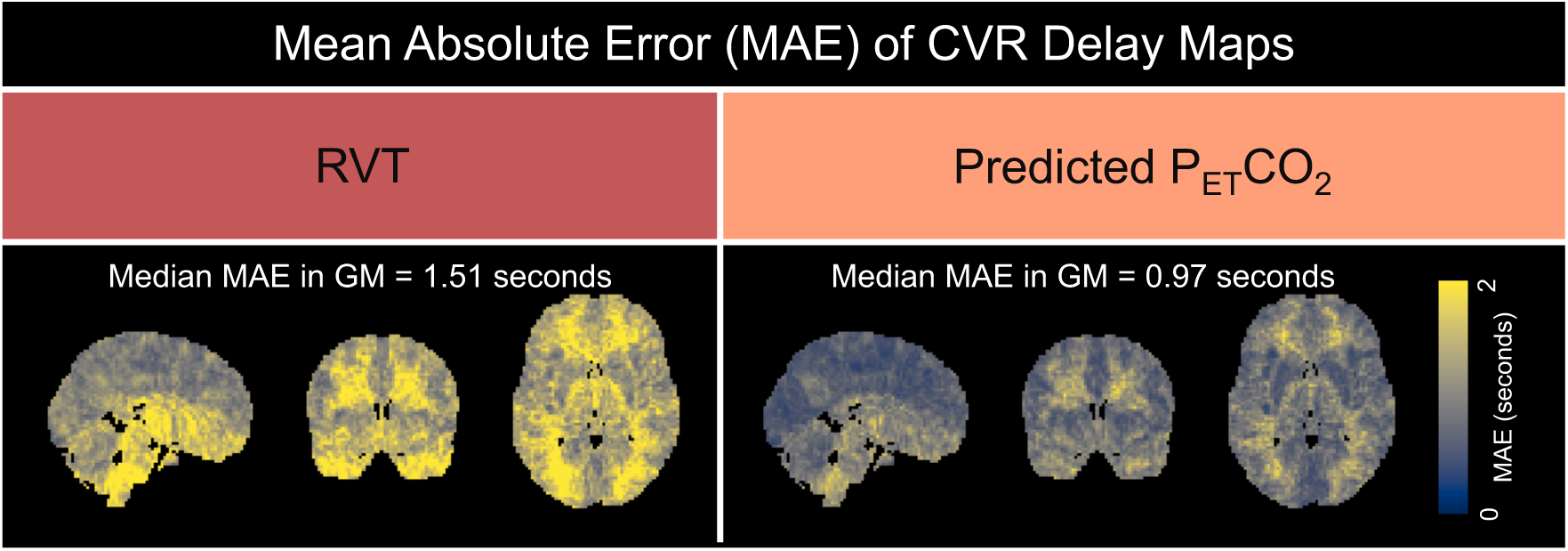
MAE maps comparing delay values generated using rescaled RVT and rescaled, predicted P_ET_CO_2_ to the ground-truth delay values generated using measured P_ET_CO_2_. The median MAE in gray matter (%BOLD/mmHg) is shown above each map.

At the scan level, within gray matter, we also evaluated the normalized correlation between the CVR amplitude values calculated using each regressor and the ground-truth amplitude values (Figure 9A). We found that CVR maps generated using predicted P_ET_CO_2_ had a significantly higher correlation to the ground-truth maps than those generated using RVT (*p*<0.05). The repeated-measures 2-way ANOVA showed that the choice of regressor and the number of breath holds used for rescaling had statistically significant main effects on the error terms (MAE and RMSE) of CVR amplitude in gray matter (*p*<0.05). Post hoc tests (Figure 9B) showed that when rescaling was performed using 1 or 2 breath holds, the MAEs and RMSEs for predicted P_ET_CO_2_ were significantly lower than those for RVT (*p*<0.05, Bonferroni corrected). When rescaling was performed using 3 breath holds, the RMSE but not the MAE was significantly lower for predicted P_ET_CO_2_ compared to RVT (*p*<0.05, Bonferroni corrected). For rescaling RVT, using 2 breath holds compared to 1 resulted in significantly decreased error terms (*p*<0.05, Bonferroni corrected), but 3 breath holds compared to 2 did not significantly change the error terms. Using more breath holds for rescaling predicted P_ET_CO_2_ did not significantly change the error terms. Effect sizes and *p*-values for all comparisons are shown in Supplementary Table S5.

**Figure 9.**
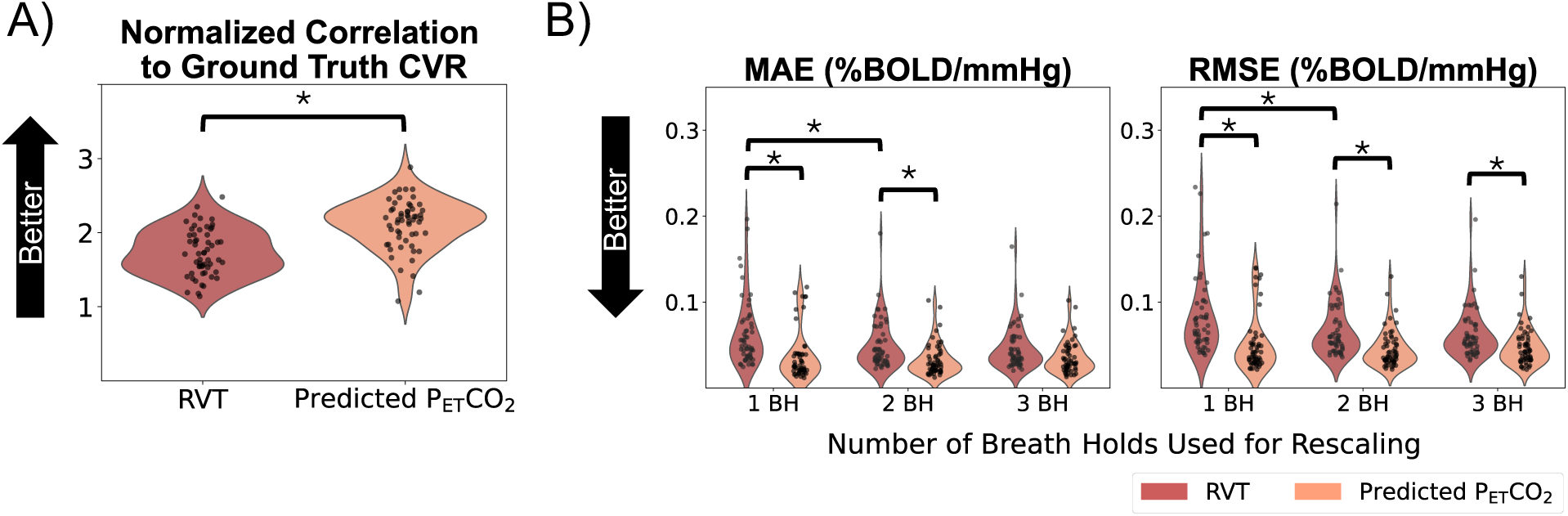
Overview of metrics comparing CVR amplitude values generated using RVT and predicted P_ET_CO_2_ to ground-truth CVR amplitude values generated using measured P_ET_CO_2_. Both Fisher’s Z values (A), which are not affected by rescaling, and error terms for each rescaling method (B) are shown. Significantly different groups are indicated by asterisks.

Lastly, we calculated Spearman rank correlations to evaluate whether the RVT and predicted P_ET_CO_2_ regressors preserve the ranking of median CVR amplitude values in gray matter across scans (Table 1). All 3 rescaling methods and both regressors resulted in CVR amplitude rankings that were significantly correlated to the ground-truth amplitude rankings. For both predicted P_ET_CO_2_ and RVT, using more breath holds for rescaling increased the Spearman rank correlation with the ground-truth amplitude values. Across the 3 rescaling methods, the amplitude rankings from rescaled, predicted P_ET_CO_2_ showed a higher Spearman correlation with the ground-truth amplitude rankings than those from rescaled RVT.

**Table 1.**
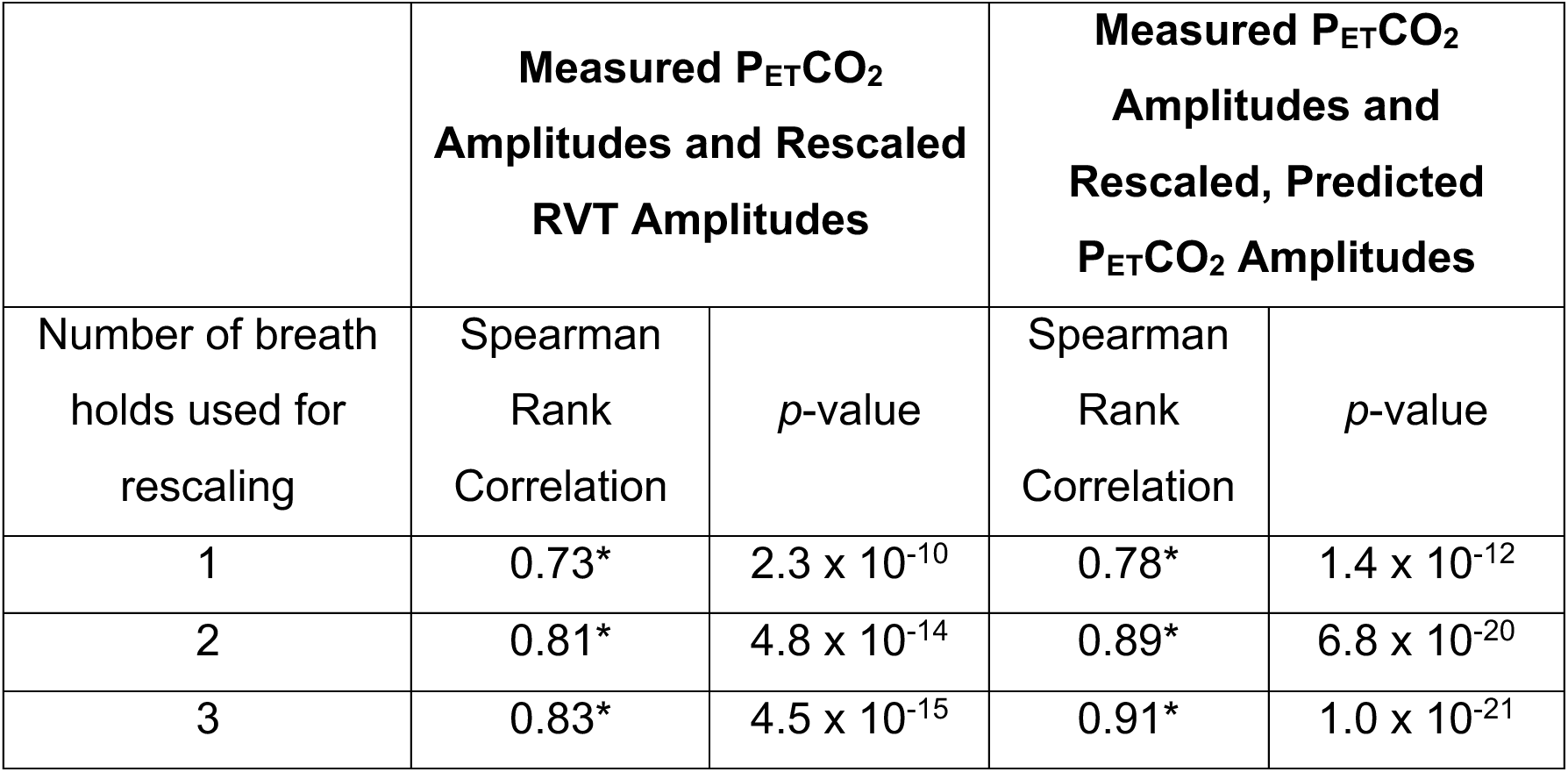
Spearman’s rank correlations and associated *p*-values comparing the median CVR amplitudes in gray matter generated using measured P_ET_CO_2_ to those generated using RVT and predicted P_ET_CO_2_ for each of the 3 rescaling methods. Asterisks indicate significant correlations (*p*<0.05).

### 3.5. CVR amplitude and delay maps for scans with low-quality breath holds

Next, we evaluated the utility of using a rescaled RVT or rescaled, predicted P_ET_CO_2_ regressor to map CVR in 2 scans with measured P_ET_CO_2_ regressors containing mostly low-quality breath holds (Figure 10). As anticipated, the amplitude maps generated using measured P_ET_CO_2_ lack the expected contrast between gray and white matter. Additionally, the measured P_ET_CO_2_ amplitude and delay maps do not appear spatially similar to a reference CVR map for the same subject from a different session with superior P_ET_CO_2_ quality. In comparison, the maps generated using rescaled RVT and rescaled, predicted P_ET_CO_2_ regressors appear more spatially similar to the reference maps, which is supported by their higher spatial correlations in gray matter to the reference maps. This suggests that rescaled RVT and rescaled, predicted P_ET_CO_2_ regressors can be used to recover reasonable maps of CVR amplitude and delay. However, while the CVR amplitude maps generated using rescaled RVT and rescaled, predicted P_ET_CO_2_ have relatively similar spatial correlations to the reference maps, the delay maps generated using predicted P_ET_CO_2_ consistently have a higher spatial correlation to the reference map than the delay maps generated using RVT.

**Figure 10.**
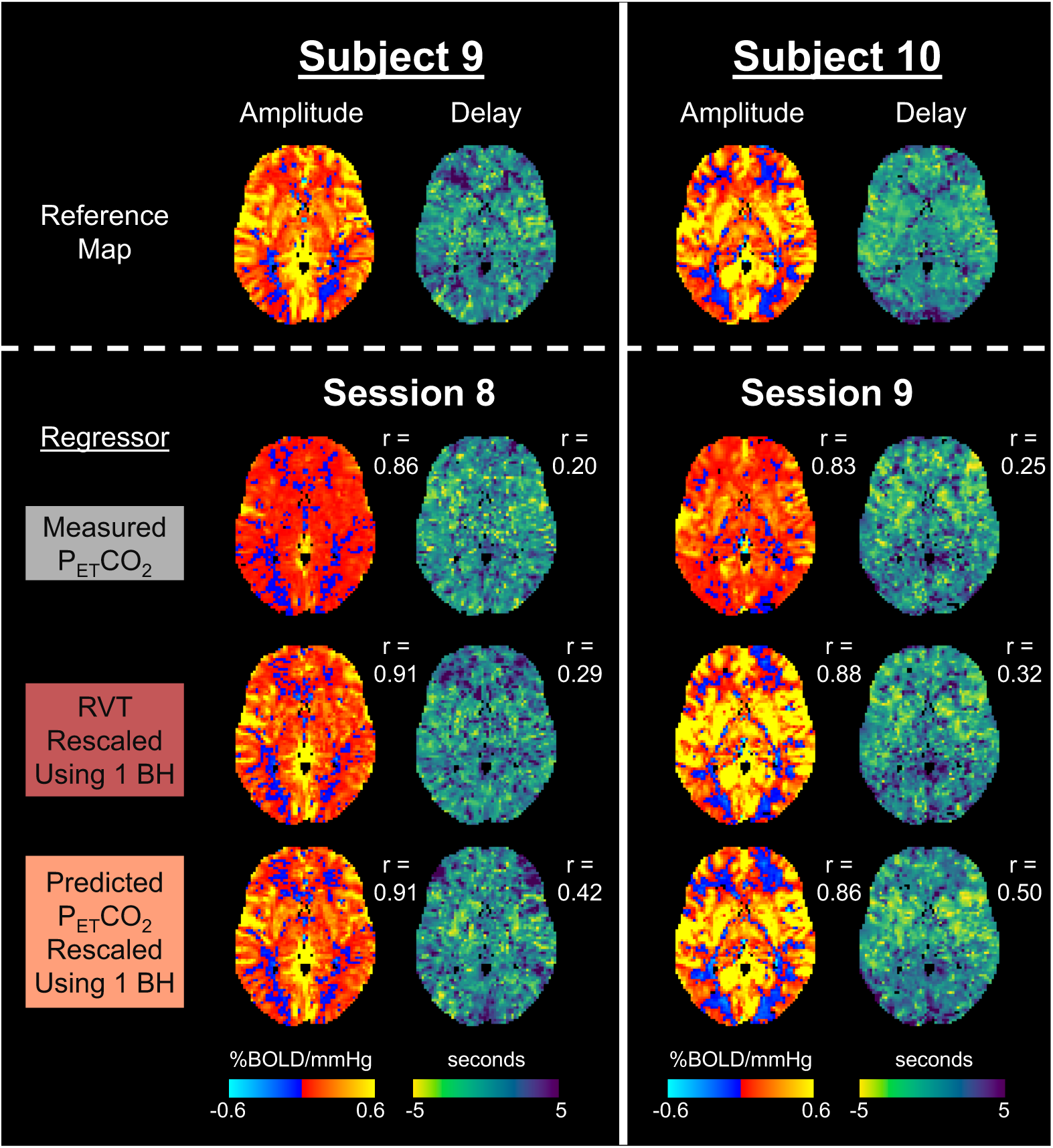
CVR amplitude and delay maps for 2 scan sessions with measured P_ET_CO_2_ timeseries containing mostly low-quality breath holds. Reference CVR amplitude and delay maps, generated using high-quality data from the same subject for a different session, are provided to assess the accuracy of maps generated using the measured P_ET_CO_2_ regressor, rescaled RVT regressor, and rescaled, predicted P_ET_CO_2_ regressor. Spatial correlations in gray matter to the reference map (r) are provided.

### 3.6. Case study in a participant with Moyamoya disease

Lastly, we evaluated the rescaled RVT and rescaled, predicted P_ET_CO_2_ regressors for mapping CVR amplitude and delay in a participant with unilateral Moyamoya disease affecting the right MCA territory. For this participant, each breath hold caused clear CO_2_ changes (average change = 6.47 mmHg) and thus we could use the CVR amplitude and delay maps generated using the measured P_ET_CO_2_ trace as a reasonable ground truth. The first breath hold caused a CO_2_ increase of 6.37 mmHg and was used for rescaling the RVT and predicted P_ET_CO_2_ regressors. As shown in Figure 11, the ground-truth CVR amplitude map does not appear to be significantly impacted by pathology. Compared to the ground-truth map, the CVR amplitude map generated using rescaled RVT shows more negative CVR values in the right hemisphere, particularly in the right MCA territory. In contrast, the amplitude map generated using rescaled, predicted P_ET_CO_2_ appears more similar to the ground-truth map, which is supported by its higher spatial correlation in gray matter. The ground-truth delay map shows that many voxels in the right hemisphere responded significantly later than voxels in the left hemisphere. Again, the delay map generated using rescaled, predicted P_ET_CO_2_ has a higher spatial correlation to the ground-truth delay map than the rescaled RVT delay map. Our finding that CVR delay, but not amplitude, is primarily impacted by Moyamoya pathology in the ground-truth maps agrees with previously reported results from a different scan of the same participant (Stickland et al., 2021).

**Figure 11.**
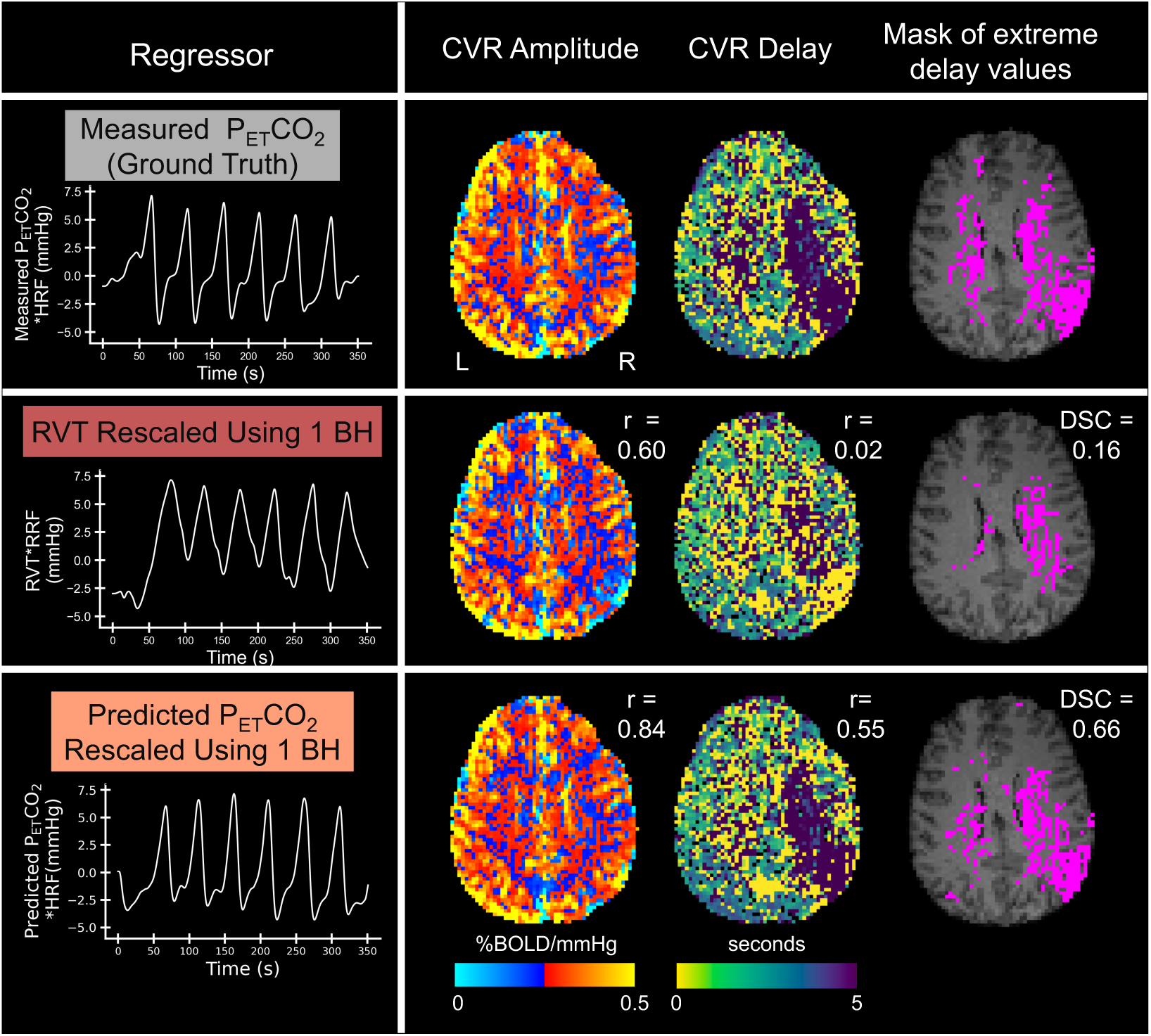
Maps of CVR amplitude, CVR delay, and extreme delay values for a participant with unilateral Moyamoya disease affecting the right middle cerebral artery. Maps were generated using 3 different regressors (from top to bottom): measured P_ET_CO_2_, rescaled RVT, and rescaled, predicted P_ET_CO_2_. For the amplitude and delay maps, spatial correlations (r) to the measured P_ET_CO_2_ (ground-truth) map in gray matter are provided. For each mask of extreme delay values, the Dice similarity coefficient (DSC) to the mask generated from the measured P_ET_CO_2_ delay map is provided.

To better understand whether RVT and predicted P_ET_CO_2_ regressors can be used to identify regions of extreme delay values, we thresholded the delay maps to isolate binary clusters with delay values greater than 10 seconds. The Dice similarity coefficient of clusters in the RVT and measured P_ET_CO_2_ delay maps was 0.16, while it was 0.66 for the clusters in predicted and measured P_ET_CO_2_ delay maps. This indicates that the CVR delay map generated using predicted P_ET_CO_2_ is more successful than RVT at characterizing this regional pathology.

## 4. Discussion

Obtaining accurate P_ET_CO_2_ data is challenging, particularly in clinical populations, which limits our ability to accurately map CVR. In this work, we explored computational methods for improving P_ET_CO_2_ data quality while maintaining units of mmHg to allow for mapping of CVR amplitude in standard units (%BOLD/mmHg) and CVR delay.

Since we observed that most participants can complete at least 1 high-quality breath-hold trial, our approach focused on leveraging high-quality measured P_ET_CO_2_ data from one or more trials to rescale 2 alternative regressors that reflect relative changes in P_ET_CO_2_ to mmHg. First, we investigated mapping CVR using a rescaled RVT regressor, which reflects changes in breathing rate and depth that cause changes in arterial CO_2_. RVT is a more feasible measure than P_ET_CO_2_ since it does not require additional task compliance from participants, but it is non-quantitative (i.e., recorded in arbitrary units). To try to better model the shape of the BOLD response to a change in arterial CO_2_ and more accurately map CVR, we also investigated whether we could predict z-normalized P_ET_CO_2_ from RVT using deep learning and rescale the predicted P_ET_CO_2_ timeseries to mmHg. Our results suggest that both rescaled RVT and rescaled, predicted P_ET_CO_2_ can be used to recover reasonable maps of CVR amplitude and delay. However, the rescaled, predicted P_ET_CO_2_ regressor is more accurate and may be more appropriate for mapping CVR in clinical populations.

### 4.1. Task compliance trends

To train and validate our P_ET_CO_2_ prediction model, we used a dataset consisting of 245 exhaled CO_2_ and respiration effort timeseries simultaneously recorded during a breath-hold task. To evaluate RVT and predicted P_ET_CO_2_ for mapping CVR, we used the publicly available EuskalIBUR dataset, consisting of fMRI data, exhaled CO_2_ timeseries, and respiration effort timeseries simultaneously recorded during a breath-hold task for 99 total scans across 10 participants. Leveraging the large sizes of these datasets, we explored general trends in breath-hold task compliance in healthy participants. It is important to note that many of the participants completed the breath hold task more than once, which may lead to higher task compliance due to learning effects. In both datasets, we found that nearly half of the CO_2_ recordings contained at least 1 low-quality breath-hold trial, indicating that CVR accuracy could be compromised in almost half of the cases. In participants with neurological diseases and in children, even lower task compliance is expected (Schlund et al., 2011; Spano et al., 2013). These results underscore the importance of developing alternative methods for mapping CVR in standard units when P_ET_CO_2_ quality is low in order to allow for CVR comparisons across subjects and scan sessions. Additionally, we found that only 4.5% and 7% of CO_2_ recordings in the training and EuskalIBUR datasets, respectively, did not contain any high-quality breath-hold trials. This finding demonstrates the overall feasibility of breath-hold tasks; while imperfect task compliance is common, only a small percentage of recordings were completely unusable. Importantly, this finding also suggests that rescaling using 1 high-quality breath hold is feasible in the vast majority of CO_2_ recordings; with coaching and real time feedback, we expect that even more recordings could contain at least 1 high-quality breath hold. For practical suggestions related to implementing breath-hold fMRI for CVR mapping, we refer readers to Appendix D.

### 4.2. Accuracy of P_ET_CO_2_ prediction

We trained a 1D FCN to predict P_ET_CO_2_ from RVT and, prior to convolution with the HRF, achieved a mean Fisher’s z-transformed correlation of 1.14 ± 0.32 with measured P_ET_CO_2_ on our held-out test set. Previously described results in the literature focused on predicting CO_2_ from respiration recordings in *resting-state* data and deriving P_ET_CO_2_ from the predicted CO_2_ timeseries (Agrawal et al., 2023); the authors achieved a mean Pearson correlation of 0.512 ± 0.269 to measured P_ET_CO_2_. For comparison to these results and to assess the benefits of using breath-hold instead of resting-state data, we calculated the mean Pearson correlation (not normalized to Fisher’s z) of measured and predicted P_ET_CO_2_ in the EuskalIBUR dataset and found that our model achieved a value of 0.787 ± 0.116. Ultimately, this high correlation supports our hypothesis that since breath holds cause large fluctuations in CO_2_, using breath-hold data may allow for more robust prediction of P_ET_CO_2_ than can be achieved with resting-state data.

Additionally, using the EuskalIBUR dataset, we found that regardless of the rescaling method used, the predicted P_ET_CO_2_ timeseries had higher normalized correlations and lower MAEs and RMSEs to measured P_ET_CO_2_ than RVT (Figure 4). This suggests that the FCN model effectively identified patterns between RVT and P_ET_CO_2_ changes, and that the predicted P_ET_CO_2_ regressor provides additional information about relative P_ET_CO_2_ changes beyond what RVT alone can provide.

### 4.3. Observations and suggestions related to rescaling P_ET_CO_2_ and RVT to mmHg

To better understand how many high-quality breath holds are required for accurate rescaling to units of mmHg, we assessed the error of predicted P_ET_CO_2_ and RVT regressors rescaled using 1, 2, and 3 breath holds relative to measured P_ET_CO_2_ (Figure 4). Rescaling predicted P_ET_CO_2_ using 2 breath holds compared to 1 breath hold significantly decreased the MAE and rescaling RVT using 3 breath holds compared to 2 breath holds significantly decreased the RMSE and MAE.

In addition to investigating whether using more breath holds for rescaling impacted the accuracy of the regressor, we also investigated how using more breath holds for rescaling impacted the actual CVR amplitude estimates (note that CVR delay is not sensitive to rescaling). By assessing the MAEs and RMSEs in gray matter for CVR amplitude values calculated using RVT (Figure 9), we found that using 2 breath holds for rescaling may be optimal (i.e., result in lower errors of CVR amplitude in gray matter) for RVT. For rescaling predicted P_ET_CO_2_, using more breath holds did not significantly decrease the MAE or RMSE in gray matter.

We did observe a small cluster of scans in which rescaling predicted P_ET_CO_2_ using a single breath hold resulted in much higher MAEs and RMSEs than the rest of the scans (as shown by the long upward tail in panel B of Figure 9). For most of these outliers, it seems that rescaling using 1 breath hold failed because the breath hold caused a smaller CO_2_ change that was not representative of the overall trace (Supplementary Figure S4A), and when rescaling was performed using 2 or 3 trials, the error decreased. When we looked at the range of CO_2_ values used for rescaling all of the datasets, we found that the range of values significantly increased as more breath holds were used for rescaling (Supplementary Figure S4B), meaning that using more breath holds decreases the likelihood that the applied rescaling underestimates the P_ET_CO_2_ changes. Therefore, if the participant completed multiple high-quality breath-hold trials, we recommend using all of them for rescaling. Adopting a more stringent threshold for high-quality trials could also prevent this issue.

If only 1 high-quality breath-hold trial is available, rescaling using 1 breath hold is sufficient in most cases. To support this, we showed that CVR amplitude and delay maps generated using RVT or predicted P_ET_CO_2_ rescaled using 1 breath hold appear highly similar to ground-truth maps (Figures 5 and 6). We also provided examples from 2 scans with low-quality measured P_ET_CO_2_ timeseries that showed that reasonable maps of CVR amplitude and delay can be recovered using RVT or predicted P_ET_CO_2_ when only 1 breath hold is used for rescaling (Figure 10). Additionally, in our case study on a participant with Moyamoya disease, we showed that when 1 breath hold is used for rescaling, predicted P_ET_CO_2_ produces CVR amplitude and delay maps that are highly similar to the ground-truth maps and sensitive to cerebrovascular pathology (Figure 11).

Another important consideration when rescaling predicted P_ET_CO_2_ or RVT regressors is defining a threshold for a high-quality breath hold; we discuss important considerations for defining this threshold in Appendix E. To ensure that at least one high-quality breath hold is collected, we recommend that researchers monitor the change in exhaled CO_2_ levels induced by each breath hold during the scan. Developing a real-time feedback tool that could automatically output whether a recently completed breath hold was high-quality would be particularly beneficial for this monitoring. If the participant does not achieve any high-quality breath holds during the task and scan time allows, researchers could ask the participants to perform additional trials.

### 4.4. Which is better: rescaled RVT or rescaled, predicted P_ET_CO_2_?

Our findings suggest that the rescaled, predicted P_ET_CO_2_ regressor produces more accurate maps of CVR amplitude and delay than rescaled RVT. Group-level MAE maps for CVR amplitude estimations showed that, across the 3 rescaling methods, predicted P_ET_CO_2_ consistently had a lower median MAE in gray matter compared to RVT (Figure 7). Additionally, group-level MAE maps for CVR delay showed that the predicted P_ET_CO_2_ regressor outperformed the RVT regressor, with a median MAE in gray matter of 0.97 compared to 1.51 seconds (Figure 8). Across the 3 rescaling methods, CVR amplitude values calculated using the predicted P_ET_CO_2_ regressor also had significantly higher normalized correlations and lower RMSEs to the ground-truth maps than those generated using the RVT regressor (Figure 9). Lastly, we found that across all 3 rescaling methods, predicted P_ET_CO_2_ better preserves the rankings of median CVR amplitudes in gray matter across scans than RVT (Table 1).

Our case study on a participant with unilateral Moyamoya disease (Figure 11) highlights the superior performance of rescaled, predicted P_ET_CO_2_ for estimating CVR amplitude and delay. Increased blood flow delays have been commonly reported in Moyamoya disease, which causes narrowing of cerebral blood vessels (Donahue et al., 2015; S. K. Kim et al., 2003; Stickland et al., 2021). In line with previous findings (Stickland et al., 2021), we found that the ground-truth CVR amplitude map was relatively unaffected by Moyamoya disease, while the ground-truth delay map showed increased delays in the right hemisphere, particularly in the vascular territory of the right middle cerebral artery, which is affected by Moyamoya disease. The amplitude and delay maps generated using rescaled, predicted P_ET_CO_2_ had higher spatial correlations to the ground-truth maps than the maps generated using rescaled RVT. In particular, the delay map generated using predicted P_ET_CO_2_ was better able to identify the region of extreme delay values compared to the RVT map. These results suggest that the predicted P_ET_CO_2_ regressor may provide the necessary sensitivity to detect impairments in CVR delay; however, given that we only had 1 patient participant, definitive conclusions cannot be made.

Additionally, more extensive research is needed to determine whether the level of accuracy associated with the rescaled, predicted P_ET_CO_2_ regressor is sufficient to identify meaningful differences in CVR amplitude across various populations. When 1 breath hold was used for rescaling, predicted P_ET_CO_2_ produced a median MAE in gray matter of 0.037 %BOLD/mmHg (Figure 7). This error is smaller than some previously reported CVR differences across populations; for example, gray matter CVR amplitude differences of 0.07 %BOLD/mmHg between young and elderly subjects have been reported (Bhogal et al., 2016). In participants with small vessel disease and traumatic brain injury, CVR amplitude differences relative to controls of 0.062-

0.079 %BOLD/mmHg and 0.042 %BOLD/mmHg, respectively, have been reported (Thrippleton et al., 2018; Bhogal et al., 2016).

### 4.5. Generalizability of P_ET_CO_2_ prediction model

To predict P_ET_CO_2_ from RVT, we trained our model using a dataset collected in our lab environment and then tested the model using the publicly available EuskalIBUR dataset. Given that our training set was relatively small, this approach helped show that our model was not overfitting to the training set and is generalizable to data collected in other research environments.

We included subject ID as an input to the model with the aim of capturing individual physiological differences, such as variations in metabolism, that influence the relationship between RVT and P_ET_CO_2_. Our model showed strong generalizability to subjects not seen during training, suggesting that subject ID may not be critical for generating accurate P_ET_CO_2_ predictions. However, including subject ID could be valuable in future studies focused on dense-mapping applications within individual subjects.

Additional work could be done to further increase the generalizability of our model. For example, to mimic participants failing to perform the trial, we collected 55 datasets in which, for each of the 10 breath holds, there was a 10% chance that the breath hold would be skipped and replaced with a period of rest. However, these 55 datasets are only a small portion of our training dataset, and our model could be improved by adding more skipped breath holds to the training dataset. With more datasets containing skipped breath holds, we could also specifically evaluate how the model predicts P_ET_CO_2_ when a breath hold is skipped. Additionally, as RVT measurements can vary with changes in belt position, incorporating datasets collected at different belt positions and from participants with different breathing styles could also increase the generalizability of the model. Since all of the datasets in the EuskalIBUR testing dataset used the same breath-hold task, we could also evaluate our model using datasets with randomized task parameters to ensure that our model is generalizable to any breath-hold task. Our in-house training dataset and the EuskalIBUR testing dataset also consisted of mostly participants in their 20s and 30s; future work could focus on collecting data in a wider age range of participants to make the model more generalizable to the broader population. Lastly, conditions such as chronic obstructive pulmonary disease and pulmonary hypertension may cause atypical relationships between ventilation (and RVT) and arterial CO_2_ and should be specifically incorporated into future model improvements (Reybrouck et al., 1998; Teopompi et al., 2013).

### 4.6. Future work

In this study, we used a 1D FCN to predict P_ET_CO_2_ from RVT, which is a relatively simple, computationally efficient approach that allows for variable-length inputs. Due to our limited training dataset size, our FCN used a discrete stopping criterion based on a fixed number of epochs. Future work will focus on integrating early stopping techniques to enhance model reliability and robustness and mitigate overfitting. Other loss functions should also be investigated in the future; we chose to use a loss function that summed the standard MSE with the MSE at the peaks scaled by 0.5. While this loss function improved peak accuracy, which is important for CVR estimation, it may have resulted in less accurate predictions between the peaks, potentially leading to less accurate rescaling of predicted P_ET_CO_2_ and RVT.

In the future, other types of models could also be investigated for predicting P_ET_CO_2_ when measured P_ET_CO_2_ quality is low. One alternative approach is using a time series forecasting model to predict low-quality segments of a P_ET_CO_2_ timeseries from high-quality segments earlier in the timeseries. In this approach, the RVT timeseries could be included as a covariate. By focusing on forecasting a part of the P_ET_CO_2_ timeseries rather than predicting the entire time series, this approach may allow the P_ET_CO_2_ predictions to be in mmHg, eliminating the need for an additional rescaling step. Another alternative model is a U-Net, which incorporates skip connections to prevent the vanishing gradients problem and has been used to successfully predict respiratory volume fluctuations from fMRI data (Bayrak et al., 2020). Simulated fMRI data from measured P_ET_CO_2_ timeseries may also be valuable in future studies for validating maps generated using rescaled, predicted P_ET_CO_2_ and rescaled RVT regressors against a known, ground-truth CVR map.

Additionally, while we showed that the predicted P_ET_CO_2_ method can be used to identify brain regions of extreme CVR delays in a single case study of an individual with unilateral Moyamoya disease, more extensive validation is needed to establish the sensitivity of the CVR amplitude and delay maps generated using rescaled, predicted P_ET_CO_2_ and rescaled RVT regressors to cerebrovascular pathology. Given the high breath-hold task compliance observed in the participant with Moyamoya disease, future investigations should target participants with low breath-hold task compliance to more thoroughly assess whether the proposed methods can recover CVR maps with the necessary sensitivity to pathology. Additionally, investigation in participants with CVR amplitude maps affected by cerebrovascular pathology is required, since CVR delay, rather than amplitude, was primarily affected in the participant with Moyamoya disease. Our ongoing research efforts include applying our methodology in participants with sub-acute and chronic stroke, to evaluate if RVT or predicted P_ET_CO_2_ remains suitable for delineating the pathological hemodynamics expected in this cohort (Krainik et al., 2005; Siegel et al., 2016).

## 5. Conclusions

We demonstrated that either an RVT or P_ET_CO_2_ regressor predicted from RVT can be rescaled using high-quality P_ET_CO_2_ data for at least one breath hold and used to model both the amplitude and delay of the CVR response to a breath-hold task. The predicted P_ET_CO_2_ regressor produces more accurate CVR amplitude and delay maps and may provide greater sensitivity to cerebrovascular pathologies. Importantly, our method (using either model) allows for CVR amplitude to be modeled in standard units (%BOLD/mmHg), facilitating CVR comparisons across subjects and scan sessions and the establishment of normative ranges of healthy CVR values. Ultimately, this work will increase the feasibility of CVR mapping in clinical settings where breath-hold task compliance may be variable.

## Data and Code Availability

Physiological data used for model training is available on OSF at https://doi.org/10.17605/OSF.IO/Y5CK4 (Clements et al., 2024). The EuskalIBUR dataset is available on OpenNeuro at doi:10.18112/openneuro.ds003192.v1.0.1 (Moia, Uruñuela, Ferrer, & Caballero-Gaudes, 2020). MRI pre-processing code is available at https://github.com/BrightLab-ANVIL/PreProc_BRAIN. Phys2cvr (Moia, Vigotsky, & Zvolanek, 2022), a publicly available Python tool, was used for computing CVR amplitude and delay maps. Additional analysis code is available at https://github.com/BrightLab-ANVIL/Clements_BHCVR-PredictedCO2.

## Author Contributions

**Rebecca G. Clements:** Conceptualization, Methodology, Software, Formal analysis, Investigation, Data curation, Writing – original draft, Writing – reviewing and editing, Visualization, Project administration. **Kristina M. Zvolanek:** Conceptualization, Methodology, Writing – reviewing and editing. **Neha A. Reddy:** Methodology, Writing – reviewing and editing. **Kimberly J. Hemmerling:** Formal Analysis, Writing – reviewing and editing. **Roza G. Bayrak:** Methodology, Writing – reviewing and editing. **Catie Chang:** Methodology, Writing – reviewing and editing. **Molly G. Bright:** Conceptualization, Methodology, Resources, Writing – review & editing, Supervision, Project administration, Funding acquisition.

## Funding

This work was supported by the National Science Foundation (DGE-2234667 to R.G.C.) and the National Institutes of Health (T32EB025766 to K.M.Z., N.A.R., and K.J.H., F31HL166079 to K.M.Z., and F31NS134222 to K.J.H.).

## Declaration of Competing Interests

The authors declare no competing financial interests.

## Acknowledgements

The authors would like to thank Chris Chin, Katie Friedman, Sara Hudson, Robbie Ng, and Kelly Tichenor in the Northwestern University Department of Physical Therapy and Human Movement Sciences for their contributions to data collection. In addition, this research was supported in part through the computational resources and staff contributions provided for the Quest high performance computing facility at Northwestern University which is jointly supported by the Office of the Provost, the Office for Research, and Northwestern University Information Technology. This research was also supported by the Center for Translational Imaging at Northwestern University.

## Appendix A. Details on physiological data collection methods for the in-house training dataset and EuskalIBUR testing dataset

<u>In-House Training Dataset and Moyamoya Dataset</u>: Exhaled CO_2_ and respiration effort were recorded using a nasal cannula connected to an ADInstruments gas analyzer and a BIOPAC respiratory belt, respectively. Signals from the gas analyzer and respiration belt were fed through PowerLab and recorded with LabChart (ADInstruments). All signals were acquired at 100 Hz.

<u>EuskalIBUR Testing Dataset</u>: During each fMRI scan, exhaled CO_2_ was measured using a nasal cannula connected to an ADInstruments gas analyzer and transferred to a BIOPAC MP150 physiological monitoring system. Respiration effort was also measured; for the first 6-7 sessions (varied between subjects), a BIOPAC respiratory effort transducer connected to a BIOPAC respiration amplifier was used, and for the remaining sessions a BIOPAC pressure pad and transducer amplifier were used. All signals were acquired at 10 kHz and down sampled to 100 Hz before any additional processing was performed.

## Appendix B: Rationale for not convolving P_ET_CO_2_ or RVT with response functions prior to training the model

P_ET_CO_2_ is often convolved with the canonical hemodynamic response function (HRF) consisting of the sum of two gamma functions (Friston et al., 1998); however, this may not be a universally optimal approach as the shape of the HRF is known to vary across subjects and brain regions (Handwerker et al., 2004). Additionally, the canonical HRF was designed to model the BOLD response to neural activity, not a CO_2_ change, and these responses have been shown to exhibit different shapes (Golestani et al., 2015). Similarly, RVT is often convolved with the respiration response function (RRF) (Birn et al., 2008), which was designed to model the average respiration-induced response function across the brain and does not account for variations in the shape of the response across subjects or brain regions. Developing a model that could predict P_ET_CO_2_ before convolution provides the flexibility to choose any response function before calculating CVR.

## Appendix C. Methods for generating voxelwise maps of CVR amplitude and delay

CVR amplitude and delay maps were generated using phys2cvr (Moia et al., 2024) using a temporal lag range of ±9 seconds in 0.3 second shift increments. For details about the lagged general linear model approach used by phys2cvr, readers are referred to Moia et al. (2020). In addition to modeling shifted variants of the P_ET_CO_2_ trace in our GLMs, we also modeled Legendre polynomials up to 4^th^ order and 6 demeaned motion parameters with each of their associated temporal derivatives. The delay maps were normalized by being recentered on the median delay in gray matter, and then the amplitude and normalized delay maps were thresholded to remove voxels with delay values at the boundary conditions (-9, -8.7, 8.7, or 9 s) since they were considered not optimized (Moia et al., 2020a). Lastly, amplitude and delay maps were registered to MNI space using the FSL 1mm MNI template resampled to 2.5mm resolution (FLIRT and FNIRT, FSL).

## Appendix D. Suggestions for implementing breath-hold fMRI for CVR mapping

To increase the likelihood that participants will successfully complete the breath-hold task, we strongly recommend allotting time before the scan for participants to practice the task. In particular, we recommend having the participant wear the nasal cannula and practice exhaling after each breath hold so that the researcher can check that the expected increase in CO_2_ is being measured and provide feedback if needed. Additionally, real-time monitoring of exhaled CO_2_ *or* the respiration belt during the scan is critical for assessing whether participants are even attempting the breath-hold task, distinct from whether the end-tidal information is successfully captured. An important caveat of using alternative regressors like RVT or predicted P_ET_CO_2_ to map CVR is that they require the participant to have attempted the breath-hold task; this means that even if the participant didn’t exhale immediately after the breath hold or accidentally breathed through their mouth, they still held their breath for most of the breath-hold periods in the task. If the participant does not seem to be attempting the breath holds at all, the researcher should stop the task and check on the participant, and, time permitting, ask the participant to redo the task.

Another important consideration is that, while respiration-belt measurements do not necessitate additional task compliance for accuracy—making them more feasible than P_ET_CO_2_ measurements—careful setup is required to properly measure respiration and ensure accurate RVT measurements. In particular, it is critical to ensure that the belt is not too loose or too tight to avoid signal saturation, which is more common in particularly large or small participants. Before the scan starts, we recommend instructing participants to take a deep breath in and out while monitoring the resulting signals. If the signals appear saturated, it is an indication that the tightness of the respiration belt should be adjusted (Bayer et al., 2024). Additionally, it is important to ensure that the respiratory belt is properly positioned to capture respiration effort, since some participants may rely on abdominal breathing as opposed to chest breathing. During data collection, we typically placed the belt approximately 3 inches above the participant’s navel, which was effective without adjustments for most participants. Additional guidelines for acquiring physiological data during fMRI scans are reviewed in detail by Bulte et al. (2017).

Our lab is currently adapting existing real-time analysis tools, originally used to give visual feedback for force targeting during motor-task fMRI (Reddy et al., 2024) to provide researchers with ongoing insight into breath-hold trial performance and signal quality during scanning. We anticipate that this approach will ensure that CVR is successfully mapped in the majority of clinical research subjects, using an efficient protocol that focuses on sufficient data quality for each individual instead of a one-size-fits-all acquisition approach.

## Appendix E. Considerations for defining the threshold for a high-quality breath hold

We determined the threshold CO_2_ increase for a high-quality breath hold by calculating the average CO_2_ increase minus 1 standard deviation across all of the breath holds in the dataset that caused a positive CO_2_ change. For the EuskalIBUR dataset, the mean CO_2_ increase across all breath holds causing a CO_2_ increase was 6.73 mmHg (mean increase – 1 standard deviation = 3.60 mmHg), and for the training dataset collected in our lab environment, the mean CO_2_ increase was 9.85 mmHg (mean increase – 1 standard deviation = 6.33 mmHg). It is surprising that the average CO_2_ increase in the training dataset was so much larger. Previously, average CO_2_ increases of approximately 9 mmHg (Tancredi & Hoge, 2013) and 13.4 mmHg (Murphy et al., 2011) in response to 20 second breath holds have been reported in healthy participants. This discrepancy could be related to sampling line lengths and vacuum settings resulting in dispersion of the exhaled gases and more mixing with room air, reducing the perceived CO_2_ changes. We suggest that at minimum, a threshold of 3.60 mmHg should be used for a 20 second breath hold to be considered high-quality; however, using a more stringent threshold may result in more accurate rescaling of predicted P_ET_CO_2_ or RVT regressors. Future work should focus on establishing guidelines for classifying breath holds as high-quality that are unique to the length of the breath hold, patient population, and perhaps even the individual participant.

## Notes

https://doi.org/10.17605/OSF.IO/Y5CK4

doi:10.18112/openneuro.ds003192.v1.0.1

### Competing Interest Statement

The authors have declared no competing interest.

### Summary of Updates

Additional details about the methods has been added to provide clarifications for readers. We have also added an ANOVA to assess the effect of the choice of regressor and the number of breath holds before running any post hoc t-tests. Additionally, we have improved the discussion section by further investigating the effect of using only 1 breath hold for rescaling and exploring potential future work in more detail.

## References

Agrawal, V., Zhong, X. Z., & Chen, J. J. (2023). Generating dynamic carbon-dioxide traces from respiration-belt recordings: Feasibility using neural networks and application in functional magnetic resonance imaging. Frontiers in Neuroimaging, 2, 1119539. 10.3389/FNIMG.2023.1119539

Bayer, J., Bayrak, R. G., Clements, R.G., Driver, I. D., Esteves, I., Figueiredo, P., Glen, D. R., Goodale, S. E., Miedema, M., Moia, S., Murphy, K., Picard, M. E., Pinto, J., Stickland, R. C., Zvolanek, K. M. (2024). Physiopy Community Guidelines, Zenodo. 10.5281/zenodo.11373723

Bayrak, R. G., Salas, J. A., Huo, Y., & Chang, C. (2020). A Deep Pattern Recognition Approach for Inferring Respiratory Volume Fluctuations from fMRI Data. International Conference on Medical Image Computing and Computer-Assisted Intervention, 12267, 428–436. 10.1007/978-3-030-59728-3_42

Bhogal, A. A., De Vis, J. B., Siero, J. C. W., Petersen, E. T., Luijten, P. R., Hendrikse, J., Philippens, M. E. P., & Hoogduin, H. (2016). The BOLD cerebrovascular reactivity response to progressive hypercapnia in young and elderly. NeuroImage, 139, 94–102. 10.1016/J.NEUROIMAGE.2016.06.010

Birn, R. M., Diamond, J. B., Smith, M. A., & Bandettini, P. A. (2006). Separating respiratory-variation-related fluctuations from neuronal-activity-related fluctuations in fMRI. NeuroImage, 31(4), 1536–1548. 10.1016/J.NEUROIMAGE.2006.02.048

Birn, R. M., Smith, M. A., Jones, T. B., & Bandettini, P. A. (2008). The respiration response function: The temporal dynamics of fMRI signal fluctuations related to changes in respiration. NeuroImage, 40(2), 644–654. 10.1016/J.NEUROIMAGE.2007.11.059

Bright, M. G., & Murphy, K. (2013). Reliable quantification of BOLD fMRI cerebrovascular reactivity despite poor breath-hold performance. NeuroImage, 83, 559–568. 10.1016/J.NEUROIMAGE.2013.07.007

Bulte, D., & Wartolowska, K. (2017). Monitoring cardiac and respiratory physiology during FMRI. Neuroimage, 154, 81–91.

Catchlove, S. J., Pipingas, A., Hughes, M. E., & Macpherson, H. (2018). Magnetic resonance imaging for assessment of cerebrovascular reactivity and its relationship to cognition: a systematic review. BMC neuroscience, 19, 1–15.

Chang, C., & Glover, G. H. (2009). Relationship between respiration, end-tidal CO2, and BOLD signals in resting-state fMRI. NeuroImage, 47(4), 1381. 10.1016/J.NEUROIMAGE.2009.04.048

Chiarelli, A. M., Villani, A., Mascali, D., Petsas, N., Biondetti, E., Caporale, A., Digiovanni, A., Grasso, E. A., Ajdinaj, P., D’Apolito, M., Rispoli, M. G., Sensi, S., Murphy, K., Pozzilli, C., Wise, R. G., & Tomassini, V. (2022). Cerebrovascular reactivity in multiple sclerosis is restored with reduced inflammation during immunomodulation. Scientific Reports 2022 *12*: 1, 12(1), 1–11. 10.1038/s41598-022-19113-8

Clements, R., Hemmerling, K. J., Reddy, N., Zvolanek, K., & Bright, M. (2024). Quantitative mapping of cerebrovascular reactivity amplitude and delay with breath-hold BOLD fMRI when end-tidal CO2 quality is low. OSF. [Dataset] 10.17605/OSF.IO/Y5CK4

Cox, R. W. (1996). AFNI: Software for Analysis and Visualization of Functional Magnetic Resonance Neuroimages. Computers and Biomedical Research, 29(3), 162–173. 10.1006/CBMR.1996.0014

Crosby, A., & Robbins, P. A. (2004). Variability in end-tidal PCO2 and blood gas values in humans. Experimental Physiology, 88(5), 603–610. 10.1113/eph8802585

Davis, T. L., Kwong, K. K., Weisskoff, R. M., & Rosen, B. R. (1998). Calibrated functional MRI: Mapping the dynamics of oxidative metabolism. Proceedings of the National Academy of Sciences of the United States of America, 95(4), 1834–1839. 10.1073/PNAS.95.4.1834/ASSET/7B6AC16D-B019-49E8-9909-63BD75C34ECF/ASSETS/GRAPHIC/PQ0384059004.JPEG

De Vis, J. B., Bhogal, A. A., Hendrikse, J., Petersen, E. T., & Siero, J. C. W. (2018). Effect sizes of BOLD CVR, resting-state signal fluctuations and time delay measures for the assessment of hemodynamic impairment in carotid occlusion patients. NeuroImage, 179, 530–539. 10.1016/J.NEUROIMAGE.2018.06.017

Donahue, M. J., Strother, M. K., Lindsey, K. P., Hocke, L. M., Tong, Y., & Frederick, B. D. (2015). Time delay processing of hypercapnic fMRI allows quantitative parameterization of cerebrovascular reactivity and blood flow delays. Journal of Cerebral Blood Flow and Metabolism, 36(10). 10.1177/0271678X15608643

DuPre, E., Salo, T., Ahmed, Z., Bandettini, P. A., Bottenhorn, K. L., Caballero-Gaudes, C., Dowdle, L. T., Gonzalez-Castillo, J., Heunis, S., Kundu, P., Laird, A. R., Markello, R., Markiewicz, C. J., Moia, S., Staden, I., Teves, J. B., Uruñuela, E., Vaziri-Pashkam, M., Whitaker, K., & Handwerker, D. A. (2021). TE-dependent analysis of multi-echo fMRI with *tedana*. Journal of Open Source Software, 6(66), 3669. 10.21105/JOSS.03669

Friston, K. J., Fletcher, P., Josephs, O., Holmes, A., Rugg, M. D., & Turner, R. (1998). Event-Related fMRI: Characterizing Differential Responses. NeuroImage, 7(1), 30–40. 10.1006/NIMG.1997.0306

Golestani, A. M., Chang, C., Kwinta, J. B., Khatamian, Y. B., & Jean Chen, J. (2015). Mapping the end-tidal CO2 response function in the resting-state BOLD fMRI signal: Spatial specificity, test–retest reliability and effect of fMRI sampling rate. NeuroImage, 104, 266–277. 10.1016/J.NEUROIMAGE.2014.10.031

Golestani, A. M., Wei, L. L., & Chen, J. J. (2016). Quantitative mapping of cerebrovascular reactivity using resting-state BOLD fMRI: Validation in healthy adults. NeuroImage, 138, 147–163. 10.1016/J.NEUROIMAGE.2016.05.025

Handwerker, D. A., Ollinger, J. M., & D’Esposito, M. (2004). Variation of BOLD hemodynamic responses across subjects and brain regions and their effects on statistical analyses. NeuroImage, 21(4), 1639–1651. 10.1016/J.NEUROIMAGE.2003.11.029

Jenkinson, M., Beckmann, C. F., Behrens, T. E. J., Woolrich, M. W., & Smith, S. M. (2012). FSL. NeuroImage, 62(2), 782–790. 10.1016/J.NEUROIMAGE.2011.09.015

Kassner, A., Winter, J. D., Poublanc, J., Mikulis, D. J., & Crawley, A. P. (2010). Blood-oxygen level dependent MRI measures of cerebrovascular reactivity using a controlled respiratory challenge: reproducibility and gender differences. Journal of Magnetic Resonance Imaging: An Official Journal of the International Society for Magnetic Resonance in Medicine, 31(2), 298–304.

Kastrup, A., Krüger, G., Neumann-Haefelin, T., & Moseley, M. E. (2001). Assessment of cerebrovascular reactivity with functional magnetic resonance imaging: comparison of CO2 and breath holding. Magnetic Resonance Imaging, 19(1), 13–20. 10.1016/S0730-725X(01)00227-2

Kim, D., Hughes, T. M., Lipford, M. E., Craft, S., Baker, L. D., Lockhart, S. N., Whitlow, C. T., Okonmah-Obazee, S. E., Hugenschmidt, C. E., Bobinski, M., & Jung, Y. (2021). Relationship Between Cerebrovascular Reactivity and Cognition Among People With Risk of Cognitive Decline. Frontiers in Physiology, 12, 645342. 10.3389/FPHYS.2021.645342/BIBTEX

Kim, S. K., Wang, K. C., Oh, C. W., Kim, I. O., Lee, D. S., Song, I. C., & Cho, B. K. (2003). Evaluation of Cerebral Hemodynamics with Perfusion MRI in Childhood Moyamoya Disease. Pediatric Neurosurgery, 38(2), 68–75. 10.1159/000068050

Kingma, D. P., & Lei Ba, J. (2015). ADAM: A Method for Stochastic Optimization. ICLR.

Krainik, A., Hund-Georgiadis, M., Zysset, S., & Von Cramon, D. Y. (2005). Regional Impairment of Cerebrovascular Reactivity and BOLD Signal in Adults After Stroke. Stroke, 36(6), 1146–1152. 10.1161/01.STR.0000166178.40973.A7

Kundu, P., Brenowitz, N. D., Voon, V., Worbe, Y., Vértes, P. E., Inati, S. J., Saad, Z. S., Bandettini, P. A., & Bullmore, E. T. (2013). Integrated strategy for improving functional connectivity mapping using multiecho fMRI. Proceedings of the National Academy of Sciences of the United States of America, 110(40), 16187–16192. 10.1073/pnas.1301725110

Kundu, P., Inati, S. J., Evans, J. W., Luh, W. M., & Bandettini, P. A. (2012). Differentiating BOLD and non-BOLD signals in fMRI time series using multi-echo EPI. NeuroImage, 60(3), 1759–1770. 10.1016/J.NEUROIMAGE.2011.12.028

Leoni, R. F., Mazzetto-Betti, K. C., Silva, A. C., Dos Santos, A. C., de Araujo, D. B., Leite, J. P., & Pontes-Neto, O. M. (2012). Assessing cerebrovascular reactivity in carotid steno-occlusive disease using MRI BOLD and ASL techniques. Radiology research and practice, 2012(1), 268483.

Leung, J., Kim, J. A., & Kassner, A. (2016). Reproducibility of Cerebrovascular Reactivity Measures in Children Using BOLD MRI. J. MAGN. RESON. IMAGING, 43, 1191–1195. 10.1002/jmri.25063

Liu, P., De Vis, J. B., & Lu, H. (2019). Cerebrovascular reactivity (CVR) MRI with CO2 challenge: A technical review. NeuroImage, 187, 104–115. 10.1016/J.NEUROIMAGE.2018.03.047

Liu, P., Hebrank, A. C., Rodrigue, K. M., Kennedy, K. M., Section, J., Park, D. C., & Lu, H. (2013). Age-related differences in memory-encoding fMRI responses after accounting for decline in vascular reactivity. NeuroImage, 78, 415–425. 10.1016/J.NEUROIMAGE.2013.04.053

Liu, P., Li, Y., Pinho, M., Park, D. C., Welch, B. G., & Lu, H. (2017). Cerebrovascular reactivity mapping without gas challenges. NeuroImage, 146, 320–326. 10.1016/J.NEUROIMAGE.2016.11.054

Lu, H., Liu, P., Yezhuvath, U., Cheng, Y., Marshall, O., & Ge, Y. (2014). MRI Mapping of Cerebrovascular Reactivity via Gas Inhalation Challenges. JoVE (Journal of Visualized Experiments), 94, e52306. 10.3791/52306

Mandell, D. M., Han, J. S., Poublanc, J., Crawley, A. P., Stainsby, J. A., Fisher, J. A., & Mikulis, D. J. (2008). Mapping cerebrovascular reactivity using blood oxygen level-dependent MRI in patients with arterial steno-occlusive disease: comparison with arterial spin labeling MRI. Stroke, 39(7), 2021–2028.

Mathieu, F., Zeiler, F. A., Ercole, A., Monteiro, M., Kamnitsas, K., Glocker, B., Whitehouse, D. P., Das, T., Smielewski, P., Czosnyka, M., Hutchinson, P. J., Newcombe, V. F. J., & Menon, D. K. (2020). Relationship between Measures of Cerebrovascular Reactivity and Intracranial Lesion Progression in Acute Traumatic Brain Injury Patients: A CENTER-TBI Study. Neurocritical Care, 37(13), 1556– 1565. 10.1089/NEU.2019.6814

Moia, S., Stickland, R. C., Ayyagari, A., Termenon, M., Caballero-Gaudes, C., & Bright, M. G. (2020a). Voxelwise optimization of hemodynamic lags to improve regional CVR estimates in breath-hold fMRI. Proceedings of the Annual International Conference of the IEEE Engineering in Medicine and Biology Society, *2020-July*, 1489–1492. 10.1109/EMBC44109.2020.9176225

Moia, S., Termenon, M., Uruñuela, E., Chen, G., Stickland, R. C., Bright, M. G., & Caballero-Gaudes, C. (2021). ICA-based denoising strategies in breath-hold induced cerebrovascular reactivity mapping with multi echo BOLD fMRI. NeuroImage, 233, 117914. 10.1016/J.NEUROIMAGE.2021.117914

Moia, S., Uruñuela, E., Ferrer, V., & Caballero-Gaudes, C. (2020b). EuskalIBUR. OpenNeuro. [Dataset] 10.18112/OPENNEURO.DS003192.V1.0.1

Moia, S., Vigotsky, A. D., & Zvolanek, K. M. (2024). phys2cvr: A tool to compute Cerebrovascular Reactivity maps and associated lag maps. 10.5281/ZENODO.7336002

Murphy, K., Harris, A. D., & Wise, R. G. (2011). Robustly measuring vascular reactivity differences with breath-hold: Normalising stimulus-evoked and resting state BOLD fMRI data. NeuroImage, 54(1), 369–379. 10.1016/J.NEUROIMAGE.2010.07.059

Murrell, C. J., Cotter, J. D., Thomas, K. N., Lucas, S. J. E., Williams, M. J. A., & Ainslie, P. N. (2013). Cerebral blood flow and cerebrovascular reactivity at rest and during sub-maximal exercise: Effect of age and 12-week exercise training. Age, 35(3), 905. 10.1007/S11357-012-9414-X

Nighoghossian, N., Berthezene, Y., Meyer, R., Cinotti, L., Adeleine, P., Philippon, B., Fremont, J.C., & Trouillas, P. (1997). Assessment of cerebrovascular reactivity by dynamic susceptibility contrast-enhanced MR imaging. Journal of the neurological sciences, 149(2), 171–176.

Papassin, J., Heck, O., Condamine, E., Pietras, J., Detante, O., & Krainik, A. (2021). Impaired cerebrovascular reactivity is associated with recurrent stroke in patients with severe intracranial arterial stenosis: A CO2 BOLD fMRI study. Journal of Neuroradiology, 48(5), 339–345. 10.1016/J.NEURAD.2020.04.005

Paszke, A., Gross, S., Massa, F., Lerer, A., Bradbury, J., Chanan, G., Killeen, T., Lin, Z., Gimelshein, N., Antiga, L., Desmaison, A., Köpf, A., Yang, E., DeVito, Z., Raison, M., Tejani, A., Chilamkurthy, S., Steiner, B., Fang, L., … Chintala, S. (2019). PyTorch: An Imperative Style, High-Performance Deep Learning Library. Advances in Neural Information Processing Systems, 32. https://arxiv.org/abs/1912.01703v1

Peirce, J. W. (2007). PsychoPy—Psychophysics software in Python. Journal of Neuroscience Methods, 162(1–2), 8–13. 10.1016/J.JNEUMETH.2006.11.017

Peng, S. L., Chen, X., Li, Y., Rodrigue, K. M., Park, D. C., & Lu, H. (2018). Age-related changes in cerebrovascular reactivity and their relationship to cognition: A four-year longitudinal study. NeuroImage, 174, 257–262. 10.1016/J.NEUROIMAGE.2018.03.033

Pinto, J., Bright, M. G., Bulte, D. P., & Figueiredo, P. (2021). Cerebrovascular Reactivity Mapping Without Gas Challenges: A Methodological Guide. Frontiers in Physiology, 11, 1711. 10.3389/FPHYS.2020.608475/BIBTEX

Reddy, N. A., Zvolanek, K. M., Moia, S., Caballero-Gaudes, C., & Bright, M. G. (2024). Denoising task-correlated head motion from motor-task fMRI data with multi-echo ICA. Imaging Neuroscience, 2, 1–30. 10.1162/IMAG_A_00057

Reybrouck, T., Mertens, L., Schulze-Neick, I., Austenat, I., Eyskens, B., Dumoulin, M., & Gewillig, M. (1998). Ventilatory inefficiency for carbon dioxide during exercise in patients with pulmonary hypertension. Clinical Physiology, 18(4), 337–344. 10.1046/J.1365-2281.1998.00109.X

Schlund, M. W., Cataldo, M. F., Siegle, G. J., Ladouceur, C. D., Silk, J. S., Forbes, E. E., McFarland, A., Iyengar, S., Dahl, R. E., & Ryan, N. D. (2011). Pediatric functional magnetic resonance neuroimaging: tactics for encouraging task compliance. Behavioral and Brain Functions, 7, 10. 10.1186/1744-9081-7-10

Siegel, J. S., Snyder, A. Z., Ramsey, L., Shulman, G. L., & Corbetta, M. (2016). The effects of hemodynamic lag on functional connectivity and behavior after stroke. Journal of Cerebral Blood Flow & Metabolism, 36(12), 2162. 10.1177/0271678X15614846

Slessarev, M., Han, J., Mardimae, A., Prisman, E., Preiss, D., Volgyesi, G., Ansel, C., Duffin, J., & Fisher, J. A. (2007). Prospective targeting and control of end-tidal CO2 and O2 concentrations. The Journal of Physiology, 581(3), 1207–1219. 10.1113/JPHYSIOL.2007.129395

Sobczyk, O., Sam, K., Mandell, D. M., Crawley, A. P., Venkatraghavan, L., McKetton, L., Poublanc, J., Duffin, J., Fisher, J. A., & Mikulis, D. J. (2020). Cerebrovascular Reactivity Assays Collateral Function in Carotid Stenosis. Frontiers in Physiology, 11, 569390. 10.3389/FPHYS.2020.01031/BIBTEX

Sobczyk, O., Sayin, E. S., Sam, K., Poublanc, J., Duffin, J., Fisher, J. A., & Mikulis, D. J. (2021). The Reproducibility of Cerebrovascular Reactivity Across MRI Scanners. Frontiers in Physiology, 12, 668662. 10.3389/FPHYS.2021.668662/BIBTEX

Spano, V. R., Mandell, D. M., Poublanc, J., Sam, K., Battisti-Charbonney, A., Pucci, O., Han, J. S., Crawley, A. P., Fisher, J. A., & Mikulis, D. J. (2013). CO2 Blood Oxygen Level–dependent MR Mapping of Cerebrovascular Reserve in a Clinical Population: Safety, Tolerability, and Technical Feasibility. https://Doi.Org/10.1148/Radiol.12112795, 266(2), 592–598. 10.1148/RADIOL.12112795

Stickland, R. C., Zvolanek, K. M., Moia, S., Ayyagari, A., Caballero-Gaudes, C., & Bright, M. G. (2021). A practical modification to a resting state fMRI protocol for improved characterization of cerebrovascular function. NeuroImage, 239, 118306. 10.1016/J.NEUROIMAGE.2021.118306

Tancredi, F. B., & Hoge, R. D. (2013). Comparison of cerebral vascular reactivity measures obtained using breath-holding and CO 2 inhalation. Journal of Cerebral Blood Flow and Metabolism, 33(7), 1066–1074. 10.1038/JCBFM.2013.48/ASSET/IMAGES/LARGE/10.1038_JCBFM.2013.48-FIG7.JPEG

Teopompi, E., Tzani, P., Aiello, M., Ramponi, S., Visca, D., Gioia, M. R., Marangio, E., Serra, W., & Chetta, A. (2013). Ventilatory Response to Carbon Dioxide Output in Patients with Chronic Heart Failure and in Patients with Chronic Obstructive Pulmonary Disease with Comparable Exercise Capacity. Respiratory Care, 59(7), 1034–1041. 10.4187/RESPCARE.02629

Thrippleton, M. J., Shi, Y., Blair, G., Hamilton, I., Waiter, G., Schwarzbauer, C., Pernet, C., Andrews, P. J. D., Marshall, I., Doubal, F., & Wardlaw, J. M. (2018). Cerebrovascular reactivity measurement in cerebral small vessel disease: Rationale and reproducibility of a protocol for MRI acquisition and image processing. International Journal of Stroke, 13(2), 195–206. 10.1177/1747493017730740/ASSET/IMAGES/LARGE/10.1177_1747493017730740-FIG4.JPEG

Tisdall, M. D., Reuter, M., Qureshi, A., Buckner, R. L., Fischl, B., & van der Kouwe, A. J. W. (2016). Prospective motion correction with volumetric navigators (vNavs) reduces the bias and variance in brain morphometry induced by subject motion. NeuroImage, 127, 11–22. 10.1016/J.NEUROIMAGE.2015.11.054

Wise, R. G., Pattinson, K. T. S., Bulte, D. P., Chiarelli, P. A., Mayhew, S. D., Balanos, G. M., O’Connor, D. F., Pragnell, T. R., Robbins, P. A., Tracey, I., & Jezzard, P. (2007). Dynamic forcing of end-tidal carbon dioxide and oxygen applied to functional magnetic resonance imaging. Journal of Cerebral Blood Flow and Metabolism, 27(8), 1521–1532. 10.1038/SJ.JCBFM.9600465/ASSET/IMAGES/LARGE/10.1038_SJ.J CBFM.9600465-FIG4.JPEG

Yezhuvath, U. S., Uh, J., Cheng, Y., Martin-Cook, K., Weiner, M., Diaz-Arrastia, R., van Osch, M., & Lu, H. (2012). Forebrain-dominant deficit in cerebrovascular reactivity in Alzheimer’s disease. Neurobiology of Aging, 33(1), 75–82. 10.1016/J.NEUROBIOLAGING.2010.02.005

Zvolanek, K. M., Moia, S., Dean, J. N., Stickland, R. C., Caballero-Gaudes, C., & Bright, M. G. (2023). Comparing end-tidal CO2, respiration volume per time (RVT), and average gray matter signal for mapping cerebrovascular reactivity amplitude and delay with breath-hold task BOLD fMRI. NeuroImage, 272, 120038. 10.1016/J.NEUROIMAGE.2023.120038

