## Supplementary Material for "Quantitative mapping of cerebrovascular reactivity amplitude and delay with breath-hold BOLD fMRI when end-tidal CO_2_ quality is low"

**Table S1.** Task timings and number of datasets for each task in the in-house training dataset

| Task Number | 1 | 2 | 3 | 4 |
|---|---|---|---|---|
| Number of datasets | 112 | 58 | 20 | 55 |
| Initial rest period duration (s) | 20 | 15 | 0 | 0 |
| Number of trials | 7 | 5 | 6 | 10 |
| Paced breathing duration (s) | 24 | 24 | 24 | [24, 30, 36] randomized with replacement |
| Breath hold duration (s) | 18 | 18 | [10, 12, 14, 16, 18, 20] randomized without replacement | [10, 11, 12, 13, 14, 15, 16, 17, 18, 19, 20] randomized with replacement, 10% chance that each breath hold is skipped and replaced with a rest period |
| Exhalation duration (s) | 2 | 3 | 2 | 2 |

| | | | | |
|---|---|---|---|---|
| Recovery duration (s) | 6 | 6 | 6 | [6, 7, 8, 9, 10, 11, 12] randomized with replacement |
| End rest period duration (s) | 30 | 15 | 0 | 0 |

**Table S2**. Correlations between measured $P_{ET}CO_2$ convolved with the HRF and RVT convolved with the HRF

| **Datasets included** | **Average Fisher's Z between measured $P_{ET}CO_2$ and RVT** |
|---|---|
| Datasets with measured $P_{ET}CO_2$ timeseries containing only high-quality breath holds | 1.04 ± 0.20 |
| Datasets with measured $P_{ET}CO_2$ timeseries containing one or more low-quality breath holds | 0.64 ± 0.27 |

**Table S3**. Error terms and hyperparameters for the 6 best performing models

| Rank | Number of Layers | Number of Epochs | Loss Function | MAE (a.u.) | RMSE (a.u.) | RMSE at the peaks (a.u.) |
|---|---|---|---|---|---|---|
| 1 | 12 | 20 | MSE(true,predicted) +0.5*MSE($true_{peaks}$,$predicted_{peaks}$) | 0.447 | 0.601 | 0.614 |
| 2 | 10 | 5 | MSE(true,predicted) | 0.409 | 0.555 | 0.674 |
| 3 | 8 | 5 | MSE(true,predicted) +0.5*MSE($true_{peaks}$,$predicted_{peaks}$) | 0.514 | 0.693 | 0.695 |
| 4 | 10 | 5 | MSE(true,predicted) +0.5*MSE($true_{peaks}$,$predicted_{peaks}$) | 0.473 | 0.669 | 0.705 |
| 5 | 10 | 15 | MSE(true,predicted) +MSE($true_{peaks}$,$predicted_{peaks}$) | 0.468 | 0.656 | 0.706 |

| | | | | | | |
|---|---|---|---|---|---|---|
| 6 | 12 | 25 | MSE(true,predicted) | 0.383 | 0.515 | 0.710 |

**Table S4.** Effect sizes and *p*-values for 2-sided paired t-tests comparing the correlations, MAEs, and RMSEs of rescaled, predicted $P_{ET}CO_2$ and rescaled RVT. Asterisks indicate significant *p*-values

| **Comparison** | ***p*-value** | **Effect Size** |
|---|---|---|
| Correlation of predicted $P_{ET}CO_2$-1BH and RVT-1BH | 0.0000* | 0.6956 |
| MAE of predicted $P_{ET}CO_2$-1BH and RVT-1BH | 0.0000* | -1.1739 |
| MAE of predicted $P_{ET}CO_2$-2BH and RVT-2BH | 0.0000* | -1.1769 |
| MAE of predicted $P_{ET}CO_2$-3BH and RVT-3BH | 0.0000* | -1.1325 |
| MAE of RVT-1BH and RVT-2BH | 0.1645 | 0.1883 |
| MAE of RVT-2BH and RVT-3BH | 0.0036* | 0.4064 |
| MAE of predicted $P_{ET}CO_2$-1BH and predicted $P_{ET}CO_2$-2BH | 0.0041* | 0.4008 |
| MAE of predicted $P_{ET}CO_2$-2BH and predicted $P_{ET}CO_2$-3BH | 0.3043 | 0.1386 |
| RMSE of predicted $P_{ET}CO_2$-1BH and RVT-1BH | 0.0000* | -1.1585 |
| RMSE of predicted $P_{ET}CO_2$-2BH and RVT-2BH | 0.0000* | -1.1259 |
| RMSE of predicted $P_{ET}CO_2$-3BH and RVT-3BH | 0.0000* | -1.0714 |
| RMSE of RVT-1BH and RVT-2BH | 0.1887 | 0.1778 |
| RMSE of RVT-2BH and RVT-3BH | 0.0042* | 0.3994 |
| RMSE of predicted $P_{ET}CO_2$-1BH and predicted $P_{ET}CO_2$-2BH | 0.0325 | 0.2932 |
| RMSE of predicted $P_{ET}CO_2$-2BH and predicted $P_{ET}CO_2$-3BH | 0.4831 | 0.0944 |

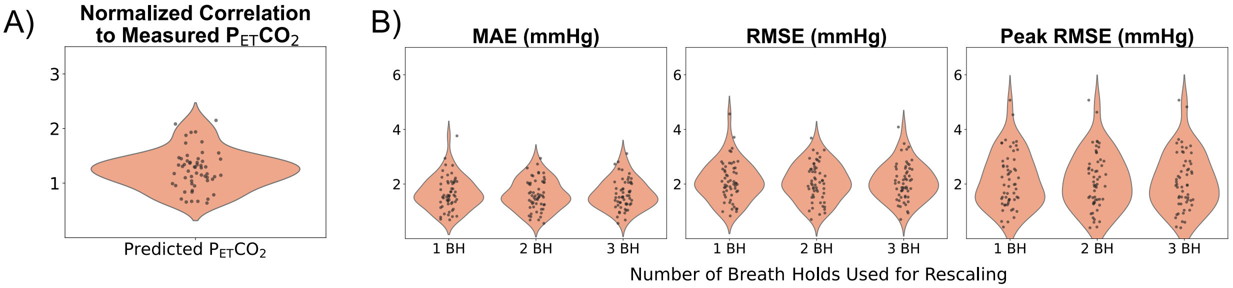


**Figure S1**. Overview of metrics comparing rescaled, predicted $P_{ET}CO_2$ to measured $P_{ET}CO_2$ in datasets in which all breath holds in the measured $P_{ET}CO_2$ timeseries were classified as high-quality. Metrics were calculated before predicted and measured $P_{ET}CO_2$ were convolved with the hemodynamic response function. Both Fisher's Z values (A), which are not affected by rescaling, and error terms for each rescaling method (B) are shown.

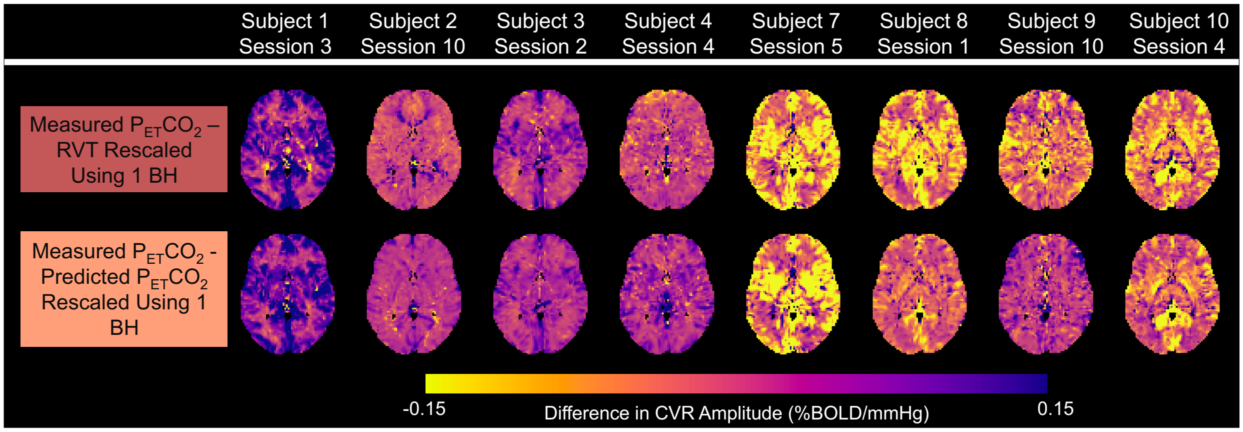


**Figure S2**. CVR amplitude difference maps for 8 subjects with measured $P_{ET}CO_2$ timeseries containing all high-quality breath holds (BHs). These maps show the differences between the ground-truth CVR amplitude values and the CVR amplitude values generated using the rescaled RVT regressor (top row), as well as the differences between the ground-truth CVR amplitude values and the CVR amplitude values generated using the rescaled, predicted $P_{ET}CO_2$ regressor (bottom row).

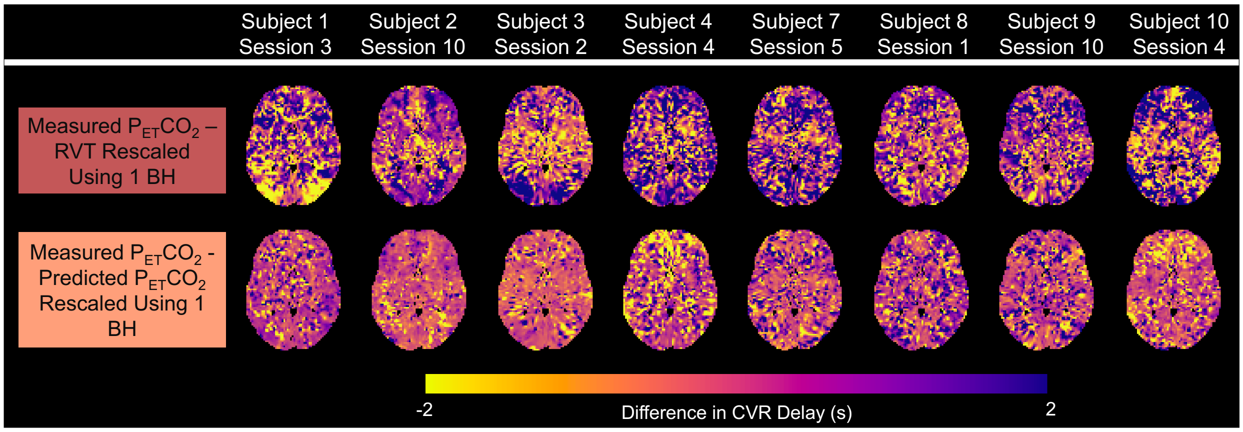


**Figure S3**. CVR delay difference maps for 8 subjects with measured $P_{ET}CO_2$ timeseries containing all high-quality breath holds (BHs). These maps show the differences between the ground-truth CVR delay values and the CVR delay values generated using the rescaled RVT regressor (top row), as well as the differences between the ground-truth CVR delay values and the CVR delay values generated using the rescaled, predicted $P_{ET}CO_2$ regressor (bottom row).

**Table S5.** Effect sizes and *p*-values for 2-sided paired t-tests comparing the correlations, MAEs, and RMSEs of CVR amplitude values in gray matter calculated using each regressor and rescaling method. Asterisks indicate significant *p*-values

| **Comparison** | ***p*-value** | **Effect Size** |
|---|---|---|
| Correlation of $CVR_{predicted\ PETCO2\text{-}1BH}$ and $CVR_{RVT\text{-}1BH}$ | 0.0000* | 1.3234 |
| MAE of $CVR_{predicted\ PETCO2\text{-}1BH}$ and $CVR_{RVT\text{-}1BH}$ | 0.0000* | -0.8604 |
| MAE of $CVR_{predicted\ PETCO2\text{-}2BH}$ and $CVR_{RVT\text{-}2BH}$ | 0.0001* | -0.5809 |
| MAE of $CVR_{predicted\ PETCO2\text{-}3BH}$ and $CVR_{RVT\text{-}3BH}$ | 0.0083 | -0.3657 |
| MAE of $CVR_{RVT\text{-}1BH}$ and $CVR_{RVT\text{-}2BH}$ | 0.0000* | 0.5952 |
| MAE of $CVR_{RVT\text{-}2BH}$ and $CVR_{RVT\text{-}3BH}$ | 0.0141 | 0.3386 |
| MAE of $CVR_{predicted\ PETCO2\text{-}1BH}$ and $CVR_{predicted\ PETCO2\text{-}2BH}$ | 0.1862 | 0.1789 |
| MAE of $CVR_{predicted\ PETCO2\text{-}2BH}$ and $CVR_{predicted\ PETCO2\text{-}3BH}$ | 0.1321 | -0.2043 |
| RMSE of $CVR_{predicted\ PETCO2\text{-}1BH}$ and $CVR_{RVT\text{-}1BH}$ | 0.0000* | -0.9413 |
| RMSE of $CVR_{predicted\ PETCO2\text{-}2BH}$ and $CVR_{RVT\text{-}2BH}$ | 0.0000* | -0.7077 |
| RMSE of $CVR_{predicted\ PETCO2\text{-}3BH}$ and $CVR_{RVT\text{-}3BH}$ | 0.0004* | -0.5063 |
| RMSE of $CVR_{RVT\text{-}1BH}$ and $CVR_{RVT\text{-}2BH}$ | 0.0000* | 0.5894 |
| RMSE of $CVR_{RVT\text{-}2BH}$ and $CVR_{RVT\text{-}3BH}$ | 0.0090 | 0.3619 |
| RMSE of $CVR_{predicted\ PETCO2\text{-}1BH}$ and $CVR_{predicted\ PETCO2\text{-}2BH}$ | 0.1165 | 0.2131 |
| RMSE of $CVR_{predicted\ PETCO2\text{-}2BH}$ and $CVR_{predicted\ PETCO2\text{-}3BH}$ | 0.2013 | -0.1728 |

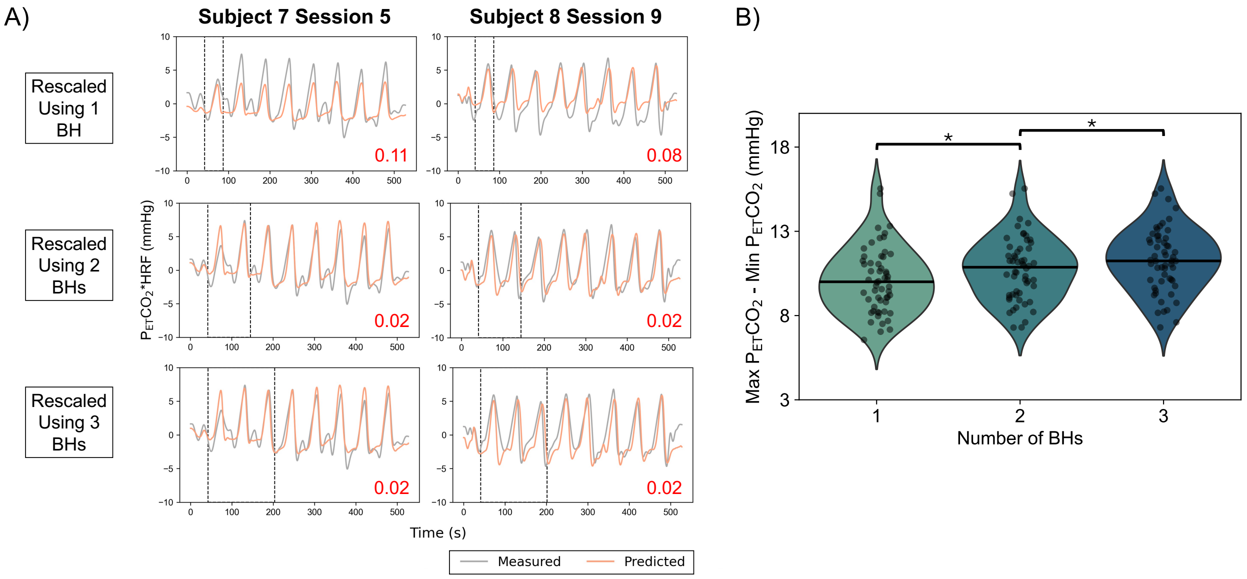

**Figure S4**. Section A shows example measured and rescaled, predicted $P_{ET}CO_2$ regressors for 2 example sessions in which rescaling using 1 breath hold (BH) was inaccurate and resulted in much higher CVR amplitude errors than rescaling using 2 or 3 BHs. Dashed lines indicate the section of measured $P_{ET}CO_2$ used for rescaling. For each rescaling method, the MAE (%BOLD/mmHg) of CVR amplitude in gray matter, calculated using the rescaled, predicted $P_{ET}CO_2$ CVR amplitude relative to the measured $P_{ET}CO_2$ CVR amplitude, is shown in red in the bottom right corner. For these example sessions, rescaling using more breath holds decreased the error of CVR amplitude. Section B shows the range of $P_{ET}CO_2$ values used for rescaling using 1, 2, and 3 breath holds for each session with all high-quality breath holds. Asterisks indicate significant differences between groups, determined using 2-sided paired *t*-tests (significance threshold $p<0.05$, with Bonferroni correction).
